# Metabolically Primed Multipotent Hematopoietic Progenitors Fuel Innate Immunity

**DOI:** 10.1101/2023.01.24.525166

**Authors:** Jason Cosgrove, Anne-Marie Lyne, Ildefonso Rodriguez, Vincent Cabeli, Cecile Conrad, Sabrina Tenreira-Bento, Emilie Tubeuf, Erica Russo, Fanny Tabarin, Yannis Belloucif, Shayda Maleki-Toyserkani, Sophie Reed, Federica Monaco, Ann Ager, Camille Lobry, Philippe Bousso, Pablo Jose Fernández-Marcos, Herve Isambert, Rafael J. Argüello, Leïla Perié

**Affiliations:** Institut Curie, Université PSL, Sorbonne Université, CNRS UMR168, Laboratoire Physico Chimie Curie, Paris, France; Metabolic Syndrome Group – BIOPROMET, Madrid Institute for Advanced Studies – IMDEA, Madrid, Spain; Inserm U944, CNRS UMR7212, Institut de Recherche Saint Louis and Université Paris-Cité, 75010 Paris, France; Dynamics of Immune Responses Unit, Institut Pasteur, Université Paris Cité, INSERM U1223, 75015 Paris, France; BMBC platform, Institut Curie, Université PSL, Sorbonne Université, CNRS UMR168, Laboratoire Physico Chimie Curie, Paris, France; Systems Immunity University Research Institute, School of Medicine, Cardiff University, Cardiff, UK; Aix Marseille Univ, CNRS, INSERM, CIML, Centre d’Immunologie de Marseille-Luminy, Marseille, France

## Abstract

Following infection, hematopoietic stem and progenitor cells (HSPCs) support immunity by increasing the rate of innate immune cell production but the metabolic cues that guide this process are unknown. To address this question, we developed MetaFate, a method to trace the metabolic expression state and developmental fate of single cells *in vivo*. Using MetaFate we identified a gene expression program of metabolic enzymes and transporters that confers differences in myeloid differentiation potential in a subset of HSPCs that express CD62L. Using single-cell metabolic profiling, we confirmed that CD62L^high^ myeloid-biased HSPCs have an increased dependency on oxidative phosphorylation and glucose metabolism. Importantly, metabolism actively regulates immune-cell production, with overexpression of the glucose-6-phosphate dehydrogenase enzyme of the pentose phosphate pathway skewing MPP output from B-lymphocytes towards the myeloid lineages, and expansion of CD62L^high^ HSPCs occurring to support emergency myelopoiesis. Collectively, our data reveal the metabolic cues that instruct innate immune cell development, highlighting a key role for the pentose phosphate pathway. More broadly, our results show that HSPC metabolism can be manipulated to alter the cellular composition of the immune system.

## Introduction

Throughout the body, stem cells need to constantly adapt the amount and type of cells that they produce to maintain tissue homeostasis, compensate for cell loss and promote tissue repair. A key challenge in the stem cell field is to understand the molecular signals that selectively differentiate stem cells into specialized cell types *in vivo*. Much emphasis has been placed on the role of transcription and growth factors in guiding lineage specification, but the role of metabolism in this context has been overlooked. Under homeostasis, hematopoiesis is the dominant biosynthetic process in the human body, generating 10^11^ cells each day^1,2^ and accounting for ~86% of daily cell turnover ^1^. Following infection, hematopoietic stem and progenitor cells (HSPCs) rapidly change the amount and the types of cells that they produce to support the immune system^3^. Despite the significant biosynthetic demands placed on HSPCs, the metabolic cues that regulate the magnitude and lineage-specificity of their cellular outputs are poorly understood.

Recently developed population-level metabolomics, genetics, and pharmacological approaches show that cellular metabolism is a key regulator of hematopoietic stem cell (HSC) function^4,5^. At the top of the hematopoietic hierarchy, HSCs have been characterized by high rates of glycolysis^6^ and low rates of protein synthesis^7^, while glutamine^8^, mitophagy and fatty acid oxidation^9,10^, vitamin A^11^, ascorbate^12^ and aspartate^13^ have been shown to regulate HSC erythroid commitment, renewal, dormancy, abundance, and reconstitution capacity respectively. As in many other stem cell systems, much work has focused on the metabolic regulation of stemness and quiescence, with relatively little focus given to the metabolic cues that guide downstream lineage specification^5,14,15^. Consequently, the metabolic processes needed to differentiate HSCs into immune cells are unknown. In addition, fate-mapping and cellular barcoding studies have shown that the downstream multipotent progenitor (MPP) compartment acts as the major source of new blood cells in native haematopoiesis^16^, and is where lineage branchpoints occur ^17,18^. Despite their functional importance, it is unclear to what extent metabolism can shape the magnitude and lineage specificity of immune cell production from MPPs.

The lack of metabolomics studies in HSPCs is due in part to the technical challenges associated with measuring metabolic processes in rare cell types. Metabolites are typically short lived - on the order of minutes^19^ and have a large structural diversity, limiting state of the art mass-spectrometry based assays to 10^4^ HSPCs^12,20^. Such limitations in sensitivity make it difficult to link relative metabolite measurements of bulk populations to the functional heterogeneity of individual HSPCs, as characterised by lineage tracing and single cell transplantation studies^17,18,21,22^. Recent advances in high-dimensional mass cytometry ^23^, flow cytometric profiling of translation^24^, genetically encoded biosensors ^25^, and *in situ* dehydrogenase assays^26^ are helping to address this challenge. However, these techniques are typically destructive in nature, making it challenging to link metabolic state to functional outcomes, particularly *in vivo*. This limitation is critical, with recent studies showing that - omics profiling should be paired with functional measurements to resolve HSPC heterogeneity ^27,28^.

To address these challenges, we developed MetaFate, an *in situ* barcoding approach to combine the metabolic gene expression state and differentiation fate of single cells *in vivo*. RNA expression has a complex association with metabolite levels, due in part to non-linear enzyme kinetics and metabolites being processed by multiple pathways^29^. RNA measurements do however provide information about the expression patterns of metabolic enzymes and transporters, and can be used to identify novel surface markers to purify functionally distinct cell-subsets for downstream metabolomics profiling. Here, using MetaFate profiling of HSPCs, we identified a gene expression program of metabolic enzymes/transporters that are associated with myeloid differentiation potential and expression of the adhesion molecule CD62L. Fluorescence based metabolic profiling assays corroborated MetaFate gene expression patterns, revealing a higher dependency on OXPHOS and glucose metabolism to fuel increased rates of protein synthesis and ATP turnover in CD62L^high^ myeloid-biased MPPs. In addition, we demonstrate that metabolism plays an active role in regulating immune cell production. MetaFate identified the pentose phosphate pathway as a metabolic signature of myeloid development. Overexpression of the glucose-6-phosphate dehydrogenase enzyme, rate limiting enzyme of the pentose phosphate pathway, limited the production of B-cells from transplanted MPPs, skewing output towards the erythromyeloid lineages. In the context of emergency myelopoiesis, the CD62L^high^ MPP compartment expands to meet increased demands for innate immune cells.

Collectively, our data bridges understanding between the fields of stem cell biology, cellular metabolism and immunology, revealing the metabolic cues that guide early innate immune cell development. Our results highlight a key role for the pentose phosphate pathway in this process and show that by manipulating lineage specific metabolic cues, it is possible to alter the specificity of regenerative processes in vivo.

## Results

### MetaFate – a lineage tracing approach to obtain fate resolved RNA expression patterns of metabolic enzymes and transporters

In this study, we hypothesise that metabolism regulates the differentiation of HSPCs *in vivo*. To address this question, we developed MetaFate, an approach that combines single cells transcriptomics with *in-situ* barcoding to provide fate-resolved expression patterns of metabolic enzymes and transporters (**Figure 1a**). For a single progenitor cell MetaFate provides 3 pieces of information: (i) gene expression data (ii) a lineage barcode (iii) the frequency at which its lineage barcode is found across differentiated cell types. Collectively, this information can be used to link metabolic enzyme/transporter RNA expression to differentiation behaviours in single cells *in vivo* (**Figure 1a**).

**Figure 1.**
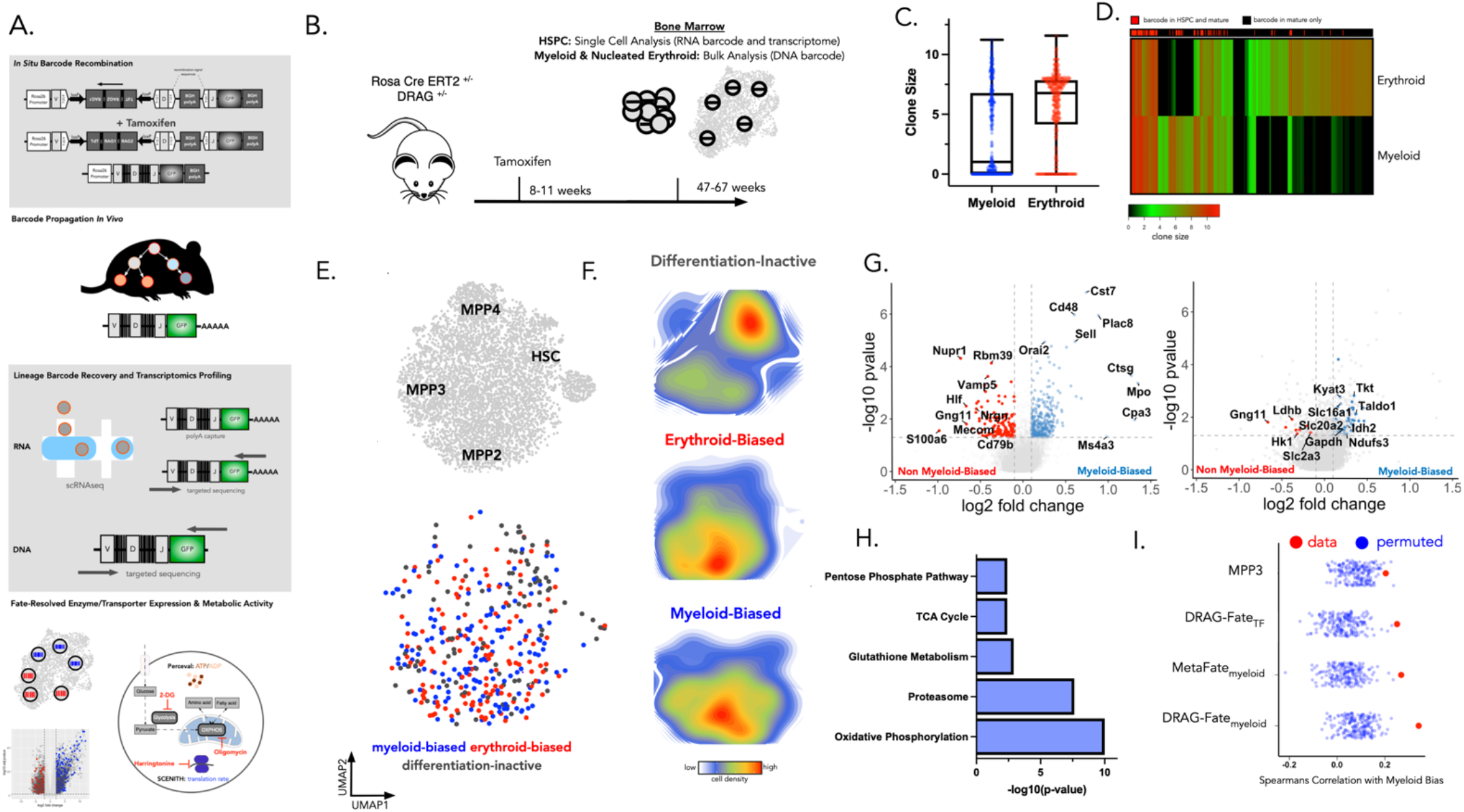
Myeloid-biased HSPCs have a distinct expression program of enzymes and transporters. (A) Overview of MetaFate and experimental set up: it consists in the induction of a lineage barcode in cells, the propagation of this barcode in vivo when cells divide and differentiate and then the recovery of the transcriptomes of HSPC and their barcode from RNA as well as the recovery of the barcode of their mature progeny by DNA. Tamoxifen injection in mice induces the recombination of the lineage barcode in situ. After division and differentiation of the barcoded cells, all offspring inheriting the barcode and a GFP tag. At the end time of the experiment, GFP-expressing HSPCs (Sca1^+^ cKit^+^), nucleated erythroid progenitors (CD44^+^ Ter119^+^) and myeloid (Cd11b^+^) cells are isolated from the bone marrow using FACS. Then bulk of mature cells were processed though nested PCR for barcode detection at the DNA level and sequenced. In parallel, HSPCs were processed through 10X scRNAseq to recover their transcriptome. Specific targeted PCR amplification were performed on the cDNA obtained by scRNAseq to recover the barcodes from the HSPC. Then, the metafate bioinformatic pipeline consolidates expression and lineage barcode data and identifies enzymes and transporters that can be targeted for functional studies, as well as surface markers to purify cell subsets for downstream metabolomics profiling. (B) Experimental timeline for induction and collection of HSPC and mature cells for metafate profiling (C) Clone sizes, number of cells per barcode, in the erythroid and myeloid lineage, the y-axis is transformed using the hyperbolic arcsin function. Each point represents a single barcode. (D) Heatmap representation of DNA barcode expression in myeloid and erythroid cells. Normalized and hyperbolic arcsin transformed cell counts (clone size) data were clustered by hierarchical clustering using Euclidian distance. Color indicates hyperbolic arcsin transformed cell counts (clone size) The top column indicates barcodes that are found both in HSPCs and mature cells (red), or barcodes found only in mature cells (black). (E) UMAP representation of the MetaFate dataset of LSK cells overlaying the positioning of known HSC and MPP subsets (top left) as well as the localisation of myeloid-biased (blue), erythroid-biased (red), and differentiation inactive (grey) progenitors based on lineage barcode (bottom left). This figure represents 4,485 Sca1^+^ cKit^+^ GFP^+^ cells (668 RNA-barcoded cells; 158 unique barcodes). (F) Density map highlighting the localization of lineage biased barcoded cells on our UMAP representation of the data. (G) Volcano plots showing differentially expressed genes between myeloid-biased barcoded cells and other (erythroid-biased and differentiation inactive) barcoded subsets. Left hand side highlights the top differentially expressed genes. All genes upregulated in myeloid-biased barcoded cells compared to erythroid and differentiation inactive-barcoded cells form a gene-signature called DRAGFate-Myeloid (271 genes) and are shown in the plot on the left hand side. The plot on the right hand side highlights differentially expressed enzymes and transporters. Downregulated and upregulated genes encoding enzymes and transporters (57 genes) are highlighted in the right hand plot in red and blue respectively. The subset of genes from the DRAGFate-myeloid signature relating to cellular metabolism form the MetaFate-myeloid signature. The signature score corresponds to the average expression values of these gene sets for each cell and is projected onto the UMAP visualisation of the data. (H) Metabolic pathways from the KEGG database that are enriched amongst genes upregulated in myeloid biased barcoded progenitors compared to erythroid and differentiation inactive barcoded cell subsets. (I) Spearmans Correlation between different transcriptomic signatures and the myeloid bias of lineage barcodes. Red points represent the correlation between signature scores and myeloid bias score, while blue points represent the correlations observed for randomised gene-sets of an equivalent size. The MPP3 signature is taken from Sommerkamp et al (2021), each point represents a different mouse. In this figure all data was taken from 5 mice from 3 independent experiments.

To barcode cells in their native environment, we use the DRAG (**D**iversity through **RAG**) *in situ* barcoding technology that allows inducible labelling of cellular lineages with heritable barcodes (**Figure 1a**, **Figure S1a**) (*Urbanus and Cosgrove et al, under revision, manuscript provided within this submission*). Importantly, DRAG barcoding is neutral with respect to hematopoietic differentiation, has a high barcode diversity, and can quantify clonal output in low cell numbers (*Urbanus and Cosgrove et al*). These key attributes make DRAG barcoding well-suited to studying the clonal dynamics of HSPCs *in vivo*. In brief, upon CRE induction by tamoxifen, the cassette between two loxP sites is inverted, causing the expression of both the RAG1 and 2 enzymes and Terminal deoxynucleotidyl transferase (TdT). This leads to the generation of a heritable barcode through the recombination of the synthetic V-, D- and J-segments, with barcode diversity being generated both by RAG-mediated nucleotide deletion and TdT-mediated N-addition (**Figure S1a**). In addition, DRAG recombination results in GFP expression, facilitating the purification of barcoded cells by fluorescence activated cell sorting (FACS). To detect barcodes in large populations of differentiated cells, we use targeted amplification of the invariant region common to all barcodes and deep sequencing of genomic DNA at the population level, as in our original protocol (*Urbanus and Cosgrove et al, in revision*). To detect barcodes and gene expression information in progenitor cells, we developed a custom targeted amplification approach using primers targeted to the invariant region of DRAG barcodes to recover barcode transcripts from 10X genomics 3’ scRNAseq libraries (*supplementary information*).

To test the metafate experimental and bioinformatics pipeline, RosaCreERT2^+/-^ DRAG^+/-^ mice were given tamoxifen injections to induce barcode recombination at 8-11 weeks of age. 47-67 weeks post-induction, barcoded HSPCs (Sca1^+^ cKit^+^ GFP^+^), Cd11b^+^ GFP^+^ Myeloid and Ter119^+^ CD44^+^ GFP^+^ nucleated erythroid cells were isolated from the bone marrow of 5 mice using fluorescence activated cell sorting (FACS) (**Figure 1b; Figure S2**). HSPCs were then processed for single cell RNA sequencing and targeted barcode amplification (materials and methods). In nucleated erythroid and mature myeloid cells, barcodes were detected from cell populations at the DNA level as described in Urbanus and Cosgrove et al. Following data pre-processing, integration and quality control (**Figure S4a-c**) transcriptomes for 4,485 cells were retained post quality control filtering and a median of 3,812 genes were detected per cell. Genes which mapped to enzymes and transporters of metabolic pathways from the KEGG, GO and REACTOME database were classified as metabolically-associated for downstream analyses (3095 genes). Following QC and filtering (materials and methods), we recovered RNA barcode information for 668 hematopoietic stem and progenitor cells (14.9% recovery rate at the RNA level), corresponding to 158 unique lineage barcodes (**table S1**). From the mature erythroid and myeloid bone marrow compartment, we recovered 381 unique barcodes at the DNA level with high consistency between technical replicates (**Figure S3**), with 97 barcodes overlapping between RNA and DNA detection. Comparison of lineage barcodes detected from either DNA or RNA showed similar barcode lengths, as well as similar insertion and deletion patterns, confirming that full barcode sequences could be accurately recovered from transcripts (**Figure S1b-d).** To compute the probability that two independent cells were labelled with the same barcode, we applied a mathematical model of DRAG barcode recombination (*Urbanus and Cosgrove et al*, materials and methods) to infer the generation probability of each barcode. Most DNA and RNA detected barcodes had a low generation probability (**Figure S1e**), with barcodes present in several mice having a higher probability than barcodes present in one mice (**Figure S1f**). Therefore many barcodes with low probability to label several cells both detected in RNA and DNA were available for lineage analysis. Collectively, these analyses demonstrate that MetaFate permits flexibility in barcode recovery at the RNA or DNA level, a comprehensive bioinformatics framework to filter spurious and probable barcodes, as well as providing a high degree of barcode diversity *in vivo*. In summary, MetaFate is a robust method to integrate *in situ* barcoding and single cell transcriptomics measurements in rare cell types.

Once we had validated the MetaFate pipeline, we sought to study the metabolic regulation of lineage commitment dynamics in HSPCs. By analysing the distribution of barcodes across the myeloid and erythroid lineages, we observed that HSPCs are highly heterogeneous in the amount (**Figure 1c**) and type (**Figure 1d**) of cells that they produced. To further characterize the functional heterogeneity of HSPCs, barcode labelled HSPCs were classified as differentiation inactive (98 cells; 61 unique barcodes) if we could not detect their barcode in any mature cell compartments^22,27,30,31^, or as erythroid-biased (143 cells; 35 barcodes), myeloid-biased (143 cells; 31 barcodes), or unbiased (284 cells; 31 unique barcodes) depending on the relative abundance of the barcode across the respective lineages. Specifically, barcodes that had more than 75% of its barcode reads in the myeloid or erythroid lineage were classified as lineage-biased (**Figure 1d-e**), or otherwise classified as unbiased. Similar results were obtained with thresholds close to 75% (**Figure S4e-g**). Extreme thresholds (90% or below 55%) impacted the amount of barcodes and the magnitude of difference in gene expression, precluding robust analysis (**Figure S4e-g**). Note that the barcode generation probability was low for most barcodes, indicating that the labeling of multiple initial cells is not accounting for the classification of the barcode in the different categories (**Figure S1g**). Within our UMAP representation of the data, the distribution of differentiation inactive clones correlated with signatures of dormant HSCs^32^, while we observed a significant overlap in the distribution of erythroid and myeloid-biased clones within MPP-associated regions of UMAP space that was not resolved by unsupervised clustering on gene expression alone (**Figure S4d**) or mapped to an existing known MPP subset (**Figure 1e-f**), suggesting that metafate revealed new biased MPP subsets to be characterized. Differential expression analysis between myeloid-biased, erythroid-biased and differentiation inactive clones identified a total of 464 differentially expressed genes associated with myeloid bias, 271 of which were upregulated in myeloid-biased clones, defining the DRAG-Fate myeloid gene signature (Figure 1e-f). Among these genes were existing markers of myeloid potential including *Mpo, Ctsg, Ms4a3* and *Cpa3*^27,33,34^, confirming that metafate can identified myeloid biased cells. Interestingly, 57/271 genes within the DRAG-Fate myeloid signature encoded enzymes and transporters from metabolic pathways of the KEGG, REACTOME and GO reference databases (**Figure 1g-h**). This subset of metabolically-associated genes, hereafter called the MetaFate myeloid signature (**Figure 1g-h**), comprises genes relating to OXPHOS (*Idh2, Idh3a, Cox7b, Ndufa4, Uqcr10*), proteostasis and ribosome biogenesis (*Hdc, Kyat3, Sec61b,Slc35b1,Psmc4*), the pentose phosphate pathway (*Tkt, Taldo1, Gpi1, Pgls*) as well as genes relating to the regulation of redox state (*Gsto1, Mgst2, Txn2,Txndc11,Gpx1*) **(Figure 1g-h).** In summary, by combining analysis of the transcriptome and the lineage barcode in the bone marrow, metafate identified a new subset of myeloid-biased MPP that upregulate specific metabolic-associated genes. These results suggest that a metabolic program associated to a lineage bias is active very early in differentiation.

Within HSPCs expression of the MetaFate and DRAGFate-myeloid gene signatures are highly correlated (**Figure S5a**), suggesting that that enzyme/transporter expression state alone could be used to predict myeloid fate in HSPCs. To compare the predictive power of metabolic-associated genes from the MetaFate myeloid signature against other signatures, we computed the Spearman’s correlation coefficient between gene signature expression scores and the myeloid bias score of HSPCs (**Figure 1i**). The DRAG fate signature consistently outperformed the MetaFate myeloid signature, suggesting that metabolism is not the only program contributing to myeloid bias (**Figure 1h**). However the MetaFate myeloid signature predicted myeloid bias to a greater extent that gene sets relating to transcription factor activity (**Figure 1i**), which are established regulators of fate choice, highlighting the importance of metabolic regulation in cell fate decisions. Furthermore, in comparison to known signatures of myeloid bias in HSPCs, the MetaFate signature had a higher correlation (rho = 0.24, p-value = 2.5 x 10^-10^) with myeloid bias compared to the existing MPP3 signature^34^ (289 genes) (rho = 0.2, p-value = 1 x 10^-7^) (**Figure 1i**). DRAG-barcode derived signatures also outperformed the MPP3 signature of myeloid bias when we performed 4-fold cross-validation analysis, to assess the sensitivity of our result to overfitting (**Figure S4h**), revealing the power of combined barcoding and transcriptome in the same cells to identify lineage bias subsets. To assess the broader predictive power of the MetaFate-myeloid signature, we quantified its expression across 3 independent published scRNAseq datasets of hematopoieitic progenitors^35–37^. In these datasets, the MetaFate-myeloid signature was upregulated in myeloid progenitors relative to other progenitor subsets (**Figure S5**). Taken together these analyses showed that the MetaFate myeloid signature, comprising only genes associated with metabolism, was a robust predictor of myeloid differentiation potential of MPPs in native hematopoiesis.

To assess whether the MetaFate myeloid expression program was maintained throughout development or was transiently expressed in Lin^-^ Sca1^+^ cKit^+^ HSPCs, we assessed enzymes and transporter gene expression patterns across different phases of myeloid development. We modelled early stages of myeloid differentiation (HSC -> MPP -> cKit^+^ restricted potential progenitors) by applying the PAGA algorithm^38^ to a published scRNAseq dataset of cKit^+^ progenitors^36^ (**Figure S6**). Using this developmental trajectory inference approach, we found that the MetaFate metabolic program was not expressed in HSCs but was heterogeneously expressed within the MPP compartment and increased as cells transition from the MPP to the cKit^+^ Sca1^-^ myeloid committed progenitor compartments. This data is consistent with previous single cell studies showing that erythroid-myeloid branching can occur before the common myeloid progenitor compartment within MPPs^18,39^

In summary, MetaFate is the first approach to map metabolic gene expression states to developmental fate in single cells in vivo. By combining expression and fate analysis, MetaFate showed that MPP’s are metabolically heterogeneous and that this heterogeneity confers differences in lineage potential. Metafate revealed a metabolic-associated gene signature that starts to be expressed in MPPs following the exit of quiescence and entry into myeloid development. This early expressed metabolic-associated program is a robust predictor of myeloid potential in MPP and is reinforced upon lineage commitment and maturation. Together this suggests that the metabolic regulation of fate decisions can occur in the earliest phases of hematopoietic development, within multipotent progenitors. To build upon this result, we next assessed to what extent the metabolic-associated gene expression patterns observed with Metafate are reflective of metabolic pathway activity in MPPs.

### CD62L^high^ Multipotent Progenitors Are Characterised by a Reduced ATP/ADP Ratio and Higher Rates of Protein Synthesis and Oxidative Phosphorylation

Given that MetaFate identified a novel myeloid-biased MPP subset with a distinct expression program of enzymes and transporters, we developed a purification strategy to isolate this subset such that we could further assess their metabolic and functional properties. Differential expression analysis between barcoded HSPCs (**Figure 1g**) highlighted *Sell*, the gene encoding the adhesion molecule CD62L, as a putative marker of cells expressing the MetaFate-myeloid expression program. We also observed significant differences in *Sell* between MetaFate^low^ and MetaFate^high^ cells (cells below and above the 75^th^ percentile of MetaFate signature expression respectively, p < 0.001) (**Figure 2a**). Flow cytometry profiling showed that CD62L is heterogeneously expressed in HSPCs (**Figure 2b-c**), with high expression in a subset of MPP3 and MPP4 cells, and little to no expression in LT-HSCs, ST-HSCs and MPP2s (**Figure 2b-c**), consistent with our MetaFate analyses. Based on these results, we selected CD62L as a marker of MetaFate^high^ myeloid-biased MPPs (**Figure 2d**).

**Figure 2.**
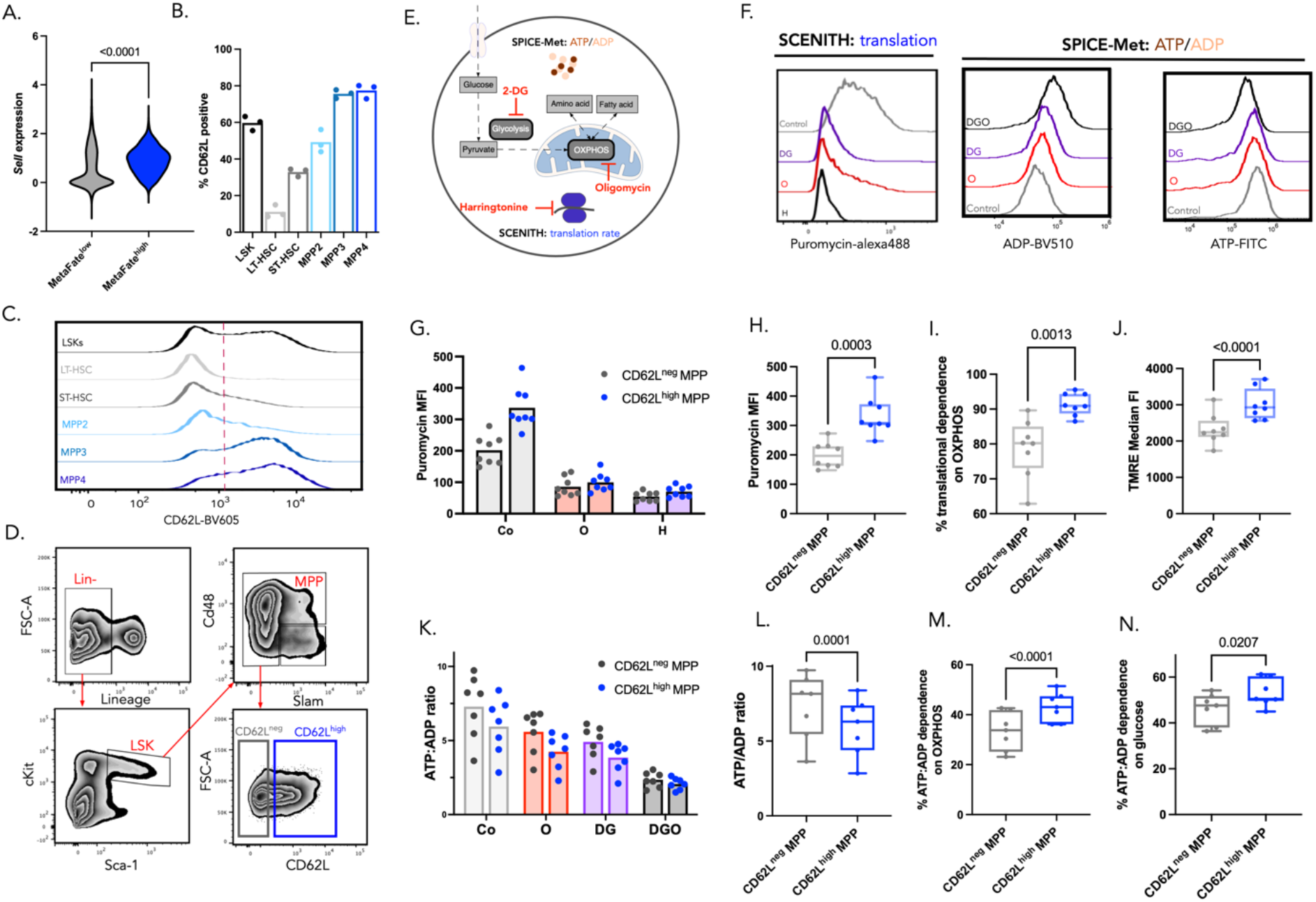
CD62L^high^ Multipotent Progenitors Are Characterised by a Reduced ATP/ADP Ratio and Higher Rates of Protein Synthesis, Oxidative Phosphorylation and Glucose Dependency: (A) Comparison of *Sell* expression between MetaFate^low/high^ expressing populations. MetaFate-low cells are defined as HSPCs in the bottom 25^th^ percentile of MetaFate signature expression. MetaFate-high cells are defined as HSPCs in the top 75^th^ percentile of MetaFate signature expression (B-C) L-selectin expression across different HSPC subsets (n = 3 mice). Gating strategy for HSPC are in figure S7a. (D) Flow cytometry gating strategy to purify L-selectin expression MPPs (E) Overview of our strategy to profile the metabolic state of different HSPC subsets. Cells are incubated in the presence of DMSO (control) or inhibitors of glycolysis (2-DG), OXPHOS (Oligomycin) or translation (Harringtonine). The scheme shows where the inhibitors are acting. Following incubation of cells with these inhibitors, we measure either translation rate (puromycin labelling), and or the percevalHR biosensor as a measure of ATP: ADP ratio. (F) Example staining profiles for SCENITH and Perceval metabolic profiling experiments on Lin-Sca1+ cKit+ bone marrow progenitors. (G) Median fluorescence intensity values for puromycin across CD62L MPP subsets and experimental conditions. Each point represents 1 mouse, data pooled from 2 independent experiments (N = 8 mice). (H) Median fluorescent intensity of puromycin labelling in CD62L^neg^ and CD62L^hi^ MPPs. (N = 8 mice, data pooled from two independent experiments). (I) Comparison of mitochondrial dependence measures calculated based on changes in puromycin labelling across control, oligomycin and harringtonine treated conditions. Formula for data transformation is provided in the materials and methods (J) Median fluorescent intensity measures for tetramethylrhodamine, ethyl ester (TMRE) labelling. N = 8 mice, data pooled from 2 experiments. Each point represents 1 mouse. Statistical comparisons were made using a paired T-test.(K) ATP:ADP measurements obtained by dividing the median fluorescent intensity of the ATP channel by the median fluorescent intensity of ADP channel. (L) Comparison of ATP:ADP ratio in CD62L^neg^ and CD62L^hi^ MPP control samples (M-N) Comparison of mitochondrial and glucose dependence measures inferred from changes in ATP:ADP ratio across control, 2-DG and oligomycin treated conditions. Formula for data transformation is provided in the materials and metods (N = 8 mice, data pooled from two independent experiments). CD62L^neg^ samples are highlighted in grey and CD62L^hi^ MPPs are highlighted in blue. Normality of the data was assessed using a Shapiro-Wilk test and statistical differences for h-j and L-N was assessed using a paired T-test. Barplots represent the mean value across all mice. Boxplots represent the median and interquartile range with whiskers extending to the minimum and maximum values.

To measure the metabolic pathway activity of CD62L^+^ MPPs, we combined two complementary fluorescence based assays (**Figure 2e,f**): (i) SCENITH (Single Cell Metabolism by Profiling Translation inhibition), a flow cytometry-based method^24^ based on profiling protein synthesis rates in response to metabolic inhibitors using flow cytometry (ii) SPICE-Met which provides a measure of cellular ATP:ADP ratio using a genetically encoded PercevalHR biosensor that can be measured using fluorescence microscopy or flow cytometry^25^. PercevalHR is composed of a mutated version of the ATP-binding bacterial protein GlnK1 and the circular permuted monomeric Venus fluorescent protein. ATP but not ADP binding to the PercevalHR causes a ratiometric shift in the probe fluorescence excitation spectrum providing a read out of ATP:ADP intracellular ratio. In these two metabolic profiling assays, cells are purified from the bone marrow of wild-type B6j (SCENITH) or Vav-iCre Perceval^fl/fl^ (SPICE-Met) mice and treated with either DMSO (control; Co), or small molecule inhibitors of glycolysis (2-deoxy-D-glucose; 2-DG), OXPHOS (Oligomycin; O) and protein synthesis (harringtonine; H). By comparing the fluorescent intensities of ATP, ADP and puromycin across different experimental conditions (**Figure 2e,f**) we can then quantify the bioenergetic state of rare cell types such as HSPCs.

To assess whether these methods could be applied to study HSPCs, we benchmarked them by comparing the metabolic profiles of HSCs (Lin^-^ cKit^+^ Cd48^-^ Slam^+^) and MPPs, for which a number of metabolic differences have already been reported^7,40,41^. Consistent with previous reports ^7,40,41^, SCENITH and SPICE-Met profiling showed that HSCs had a higher glycolytic capacity and lower protein synthesis rate than MPPs (**Figure S7b-d**), confirming that our approach can be successfully applied to study other hematopoietic progenitor subsets. To assess whether the MetaFate-myeloid signature was reflective of differences in metabolic pathway activity, we then compared the metabolic profiles of CD62L^neg^ and CD62L^high^ MPPs (Lin^-^ cKit^+^ Sca1^+^) (**Figure 2g,k**). SCENITH profiling showed that CD62L^high^ MPPs have a significantly higher rate of protein synthesis than CD62L^neg^ MPPs (p < 0.001) and their translation rates are highly sensitive to oligomycin treatment (p = 0.001) (**Figure 2h-i**). Similar results were obtained when we measured the uptake of mitochondrial membrane potential TMRE dye in the different MPP subsets (p < 0.001) **(Figure 2j)**. Using SPICE-Met, we observed that CD62L^high^ MPPs had a lower ATP/ADP ratio (p < 0.001) than CD62L^neg^ MPPs **(Figure 2k,l)** and a higher OXPHOS-dependence (p < 0.001) **(Figure 2m)**, corroborating results from SCENITH. To understand if the relationship between translation rates and OXPHOS was entirely glucose dependent, or to what extent the breakdown of fatty and amino acids via the TCA cycle was also involved, we measured ATP:ADP ratios of MPPs following inhibition of glucose metabolism using 2-DG. In this analysis, ATP:ADP ratios **(Figure 2n),** were more sensitive to glucose inhibition in CD62L^high^ MPPs compared CD62L^neg^ MPPs (p = 0.02), suggesting that the higher rates of translation observed in CD62L^high^ MPPs are typically fuelled by glucose, rather than through the oxidation of fatty and amino -acids.

Consistent with our MetaFate analyses, CD62L^high^ MPPs have distinct metabolic properties compared to other multipotent progenitors. Specifically we observed higher rates of ATP turnover and protein synthesis, with metabolic demands fuelled by increased glucose catabolism in the mitochondria, rather than by increasing the rate of fatty and amino acid oxidation. Our results suggest that metabolic remodelling accompanies the earliest phases of hematopoietic lineage specification, and so we next investigated whether metabolism plays an active or a passive role in the decision making process.

### The Pentose Phosphate Pathway Actively Regulates Immune Cell Production

MetaFate analyses and metabolic profiling highlighted a number of metabolic pathways that are associated with early myeloid development. This led us to hypothesise that manipulating metabolic processes within MPPs could influence the rate of immune cell production. To better discriminate between pathways that have an active versus a passive role in myelopoiesis, we applied the MIIC causal network reconstruction algorithm^42^ to enzyme and transporter expression data obtained from bulk RNA sequencing samples across the entire hematopoietic system ^43^ (**Figure S8**). MIIC is an information theoretic method which learns graphical models from observational data, including the effects of unobserved latent variables^42,44^. MIIC network reconstruction predicted that in myeloid cells, enzymes of the PPP (*Taldo1, G6pdx*) are strongly associated with redox state (*Gsr, Mgst1, Mgst2*), glucose metabolism (*Hk2, Pkm*), NADPH-oxidase activity (*Ncf1*) and lipid metabolism (*Scarb1, Abcd1,Acer3*), and collectively, these enzymes contribute to myeloid lineage specification. This prediction was consistent with MetaFate results, which showed that enzymes associated with the PPP are upregulated in myeloid-biased MPPs (**Figure 1h**). Together, this led to the hypothesis that manipulating the PPP in MPPs could be a strategy to regulate the dynamics of immune cell production.

To test this hypothesis, we use a murine model where Glucose-6-Phosphate-Dehydrogenase (G6PD), the rate limiting enzyme of the PPP, is overexpressed^45^. In this G6PD overexpression system a large genomic fragment (20.1Kb) of the entire human G6PD gene, including upstream and downstream regulatory sequences was inserted into the genome of a transgenic mouse line (G6PD-Tg). In G6PD-Tg mice, G6PD expression is increased 2-fold at the RNA level compared to WT littermate controls, a phenotype associated with increased G6PD enzyme activity and NADPH production rates^45^. To assess the functional consequences of G6PD overexpression on MPP differentiation, we transplanted G6PD-tg and WT MPPs and quantified their differentiation patterns *in vivo* using a lentiviral cellular barcoding approach (**Figure 3a**). A single cell lineage tracing approach such as lentiviral barcoding allows to follow the fate of heterogenous cells like MPPs. MPPs were purified from the bone marrow of G6PD-Tg mice or WT littermate controls and infected with the LG2.2 lentiviral barcode library as previously described^46^. Cells were then transplanted into sub-lethally irradiated (6Gy) recipients and left to engraft, divide and differentiate. At day 21 after transplantation, the timepoint where myeloid production from transplanted MPPs peaks^47^, barcoded (GFP^+^) erythroblasts (E; Ter119^+^ CD44^+^), myeloid cells (M; Ter119^-^ CD19^-^ CD11b^+^), and B-cells (B; Ter119^-^ CD11b-CD19^+^) were sorted from the bone marrow and their barcode identity was assessed through PCR and deep sequencing from their bulk DNA (**Figure 3a and S9a**).

**Figure 3:**
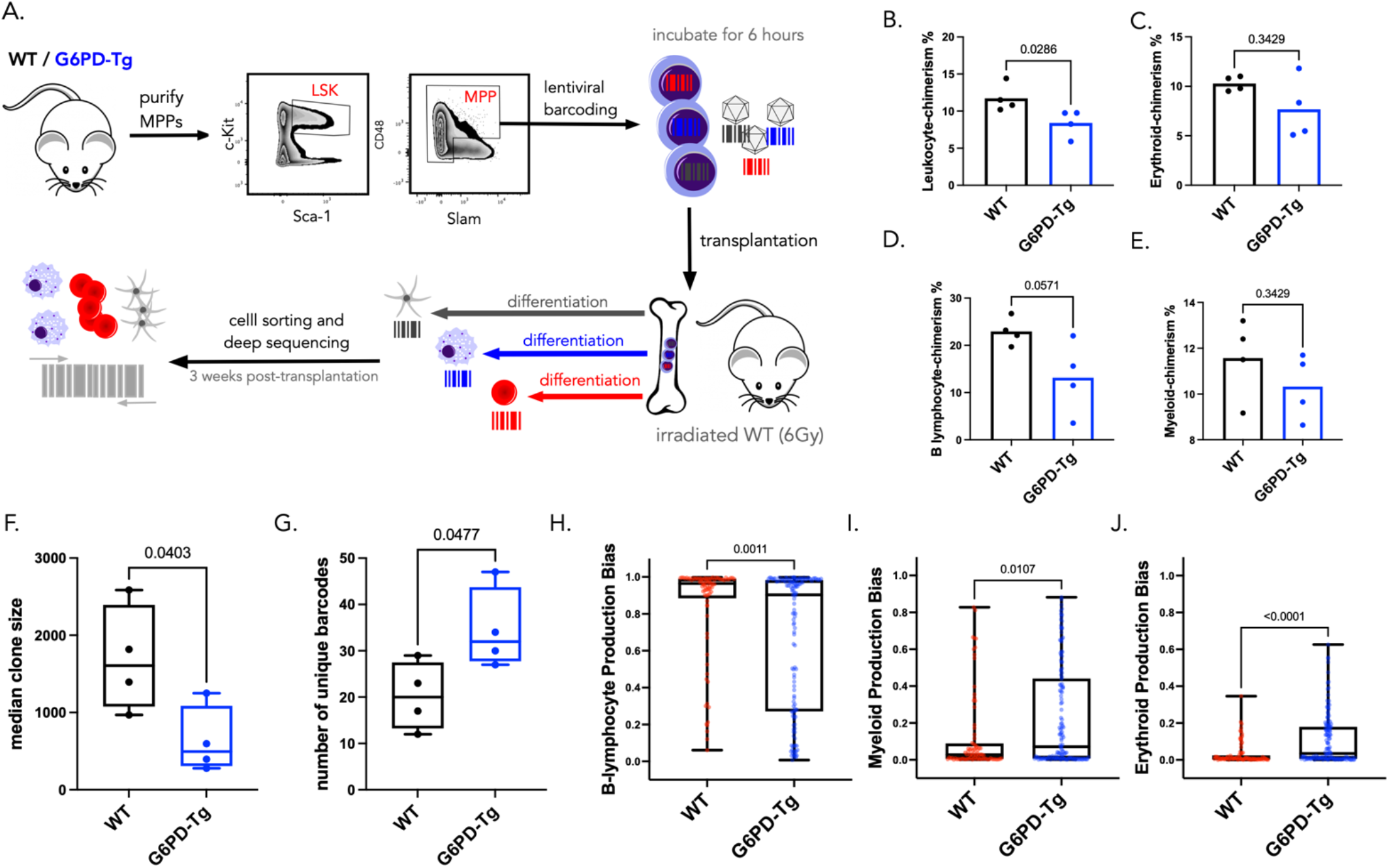
Upregulation of Glucose-6-Phosphate-Dehydrogenase in MPPs Inhibits B-Lymphopoiesis Post-Transplantation. (A) Overview of the lentiviral barcoding experiment. MPPs (say phenotype) were purified from the bone marrow of WT or G6PD-Tg mice by FACS and were infected with the LG2.2 lentiviral barcoding library. 6 hours later transduced MPPs were injected I.V. into 6Gy irradiated WT recipients. 3 weeks post-transplantation bone marrow was harvested and cells were sorted and their bulk DNA was processed for barcode detection through nested PCR and sequencing. (n = 4 mice per condition). (B-E) % chimerism quantified by measuring the proportion of GFP cells relative to total live cell numbers in the respective lineage compartments by flow cytometry for the WT (black) and G6PD-Tg (blue) transplanted MPPs. Each point represents a single mouse with N=4 mice per experimental condition. Pairwise comparisons were made using a Mann-Whitney test. (F) Median clone sizes for the top B-cell producing barcodes (the top *n* barcodes defined as contributing to 95% of all read counts for the B-cell lineage) from WT (black) and G6PD-Tg (blue) transplanted MPPs. Each point represents a single mouse. Pairwise comparisons are made using a Student’s T-test. (G) Barcode diversity as defined by the number of unique barcode for B-cell producing barcodes from WT (black) and G6PD-Tg (blue) MPPs, each point represents a single mouse. Pairwise comparisons are made using a Student’s T-test (H-J) The lineage bias of each B-cell producing barcode is shown per lineage and per experimental condition. The bias is calculated by computing the frequency of barcode *i* in lineage *j*, and then comparing the relative frequencies of each barcode across all lineages. Each point represents a single barcode (81 WT barcodes (red), 138 G6PDtg barcodes (blue)) with data pooled from 4 mice per condition. Pairwise comparisons were made using a Mann-Whitney test. Boxplots represent the median and interquartile range with whiskers extending to the minimum and maximum values.

Analysis of chimerism post-transplantation using the proportion of GFP^+^ barcoded cells in the erythroid, myeloid, and B-cell compartments showed a significant decrease in GFP^+^ cells in G6PD-Tg derived leukocytes relative to WT controls, with a trending decrease in B-cell chimerism (p = 0.057) but no change in myeloid chimerism (p = 0.34) (**Figure 3b-e**). This result show that over-expression of a myeloid associated enzyme, G6PD lead to impaired B-lymphopoiesis. To better assess this phenotype, we analysed the distribution of lentiviral barcodes amongst mature cell types. For G6PD-Tg and WT samples, we detected similar number of sequencing reads (**Figure S9b**) as well as a high consistency between PCR duplicates (**Figure S9c-d**) and very little sharing of barcodes between mice (**Figure S9e**). Following these QC steps, we analysed the diversity, clone size distributions, and lineage bias of G6PD-Tg vs WT barcoded MPPs to understand the progenitor dynamics that gave rise to reduced leukocyte chimerism. Consistent with chimerism measurements, we observed no significant differences in the diversity, clone sizes, and bias of the erythroid- and myeloid-producing barcoded MPPs between the G6PD-Tg and control group (**Figure S10c-h**). Within the B-cell lineage, the cumulative barcode read distribution showed that only a small fraction (11.9 ± 2.5% of WT and 21 ± 9% of G6PD-Tg) of transplanted cells give rise to 95% of all donor-derived B cells **(Figure S10b),** contrary to the other two lineages (**Figure S10c-h**). Focusing on these B-cell producing MPP barcodes, we observed a 2.7-fold reduction in the number of B-cells derived from each barcoded MPP in the G6PD-Tg group compared to WT **(Figure 3f)**, with a median clone size of 1692 ± 689 cells for WT B-cell barcodes, but only 630 ± 434 cells for G6PD-Tg B-cell barcodes **(Figure 3f, Figure S10i).** This reduced amount of lymphoid cells produced per MPP due to G6PD overexpression was then partly compensated at the population level by an increase the total number of MPPs producing B-cells in the G6PD-Tg group compared to WT (**Figure 3g**). Overall per individual MPP, G6PD over-expression led to a net-skewing of cell production towards the erythro-myeloid lineages at the expense of the lymphoid lineage (**Figure 3h-j**). In summary, by combining targeted genetics and cellular barcoding approaches, we showed with single cell resolution that overexpression of the pentose phosphate pathway, a key pathway from our metafate myeloid derived gene signature, limits B-cell production *in vivo*. This result confirmed our hypothesis that manipulating metabolic processes within MPPs can regulate the dynamics of immune cell production *in vivo*.

### CD62L^high^ MPPs Fuel Emergency Myelopoiesis during acute infection and bone marrow transplantation

Hematopoiesis is a highly dynamic system, and must adapt to meet changing requirements for blood and immune cells. We hypothesised that the metabolically-primed myeloid-biased multipotent progenitors we identified via MetaFate, could play a significant role in emergency myelopoiesis where the rate of myeloid cell production increases significantly^48^.

To assess the role of CD62L^high^ MPPs in infection we first used an LPS challenge model in which mice are given 35μg of LPS at 0 and 48 hours, and bone marrow samples are processed for flow cytometry analysis at 72 hours^49^ (**Figure 4a**). In this established model of emergency myelopoiesis^49^, we observe increased CD62L expression in MPPs (**Figure 4b**) and an increased proportion of CD62L^high^ MPPs (Figure 4c). This change correlated with large increases in both the GMP (cKit^+^ Sca1^-^ CD16/32^+^ CD34^-^) and myeloid (Cd11b^+^) compartments of the bone marrow (**Figure 4c**). To assess whether the CD62L^high^ HSPC expansion can also occur following infection with a live pathogen, we re-analysed a scRNAseq dataset of cKit^+^ progenitors from WT mice, or mice infected with *Plasmodium Berghei* 7 days post infection^50^ (**Figure 4d**). In this setting, our MetaFate-myeloid expression program gene signature was significantly increased in HSPCs relative to control cells (**Figure 4f,g**). Additionally, we saw a concomitant increase in the number of cells which expressed *Sell* – the gene encoding CD62L **(Figure 4e,h).** Taken together, our results show that the CD62L multipotent progenitor compartment plays a key role in supporting immune responses by producing innate immune cells.

**Figure 4:**
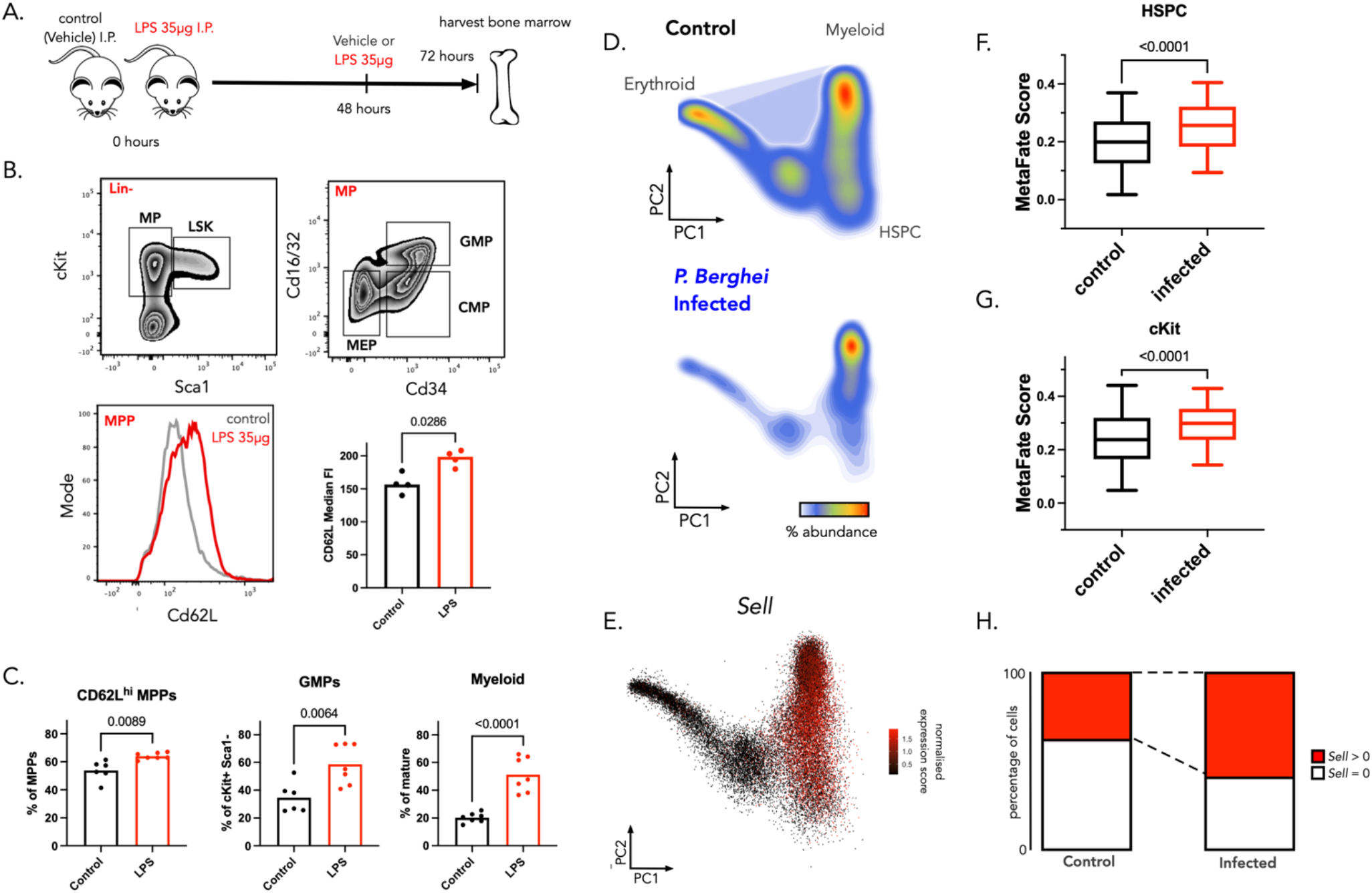
The CD62L^hi^ MPP compartment expands to fuel emergency myelopoiesis during acute infection. (A) Overview of the LPS challenge model. 18 week old B6j Mice were injected with LPS (35ug/mouse) I.P. at 0 hours and 48 hours. At 72 hours bone marrow cells were harvested and analysed by flow cytometry. (B) Gating strategy for analysis of the progenitors in the bone marrow from a representative mouse and median fluorescence intensity of CD62L expression in multipotent progenitors in control and LPS treated mice. Each point is a mouse, N= 4 mice. Statistical significance was assessed using a Mann-Whitney test(C) Quantification of the porcentage of the different cell subsets in control (black) and LPS treated (red) mice. Normality of the data was assessed using a Shapiro-Wilk test and significance was assessed using a T-test. N = 13 mice and data was pooled from 2 independent experiments. Barplots represent the mean expression value and each point represents a different mouse. (D) scRNAseq data reanalysed from Haltalli et al (2020), where mice were treated with vehicle control or *P. berghei*. 7 days post infection cKit+ progenitors were purified from the bone marrow of each group and processed for scRNAseq profiling. Data are represented using a density projection of the cell abundances on to a PCA embedding of the data. Cell type annotations of the data were taken from the original publication. Control samples have 14193 cells and infected samples have 13905 cells. (E) *Sell* (gene encoding CD62L) normalised gene expression projected onto the PCA embedding of the data. (F) Boxplot showing MetaFate signature expression score in HSPCs (defined as primitive HSPCs in the original article) from control (black) and infected (red) mice. Pairwise comparisons were made using a Students T-test. Boxplot showing mean and sd over cells? (G) Same as F but for all cKit+ hematopoietic progenitors from control and infected mice. (H) The proportion of cells in control and infected mice that have non-zero expression of *Sell*. Boxplots represent the interquartile range and median values and whiskers represent the 5^th^ and 95^th^ percentile of the data.

In the context of bone marrow transplantation, the immune system must be restored following conditioning protocols to avoid life-threatening complications^51^. To assess whether CD62L^high^ MPPs preferentially reconstitute the myeloid compartment following bone marrow transplantation, we purified CD62L^high^ and CD62L^neg^ MPPs by FACS and transplanted them into irradiated recipient mice. However the use of the MEL-14 anti-CD62L antibody clone to sort cells and transplantation led to much poorer engraftment of CD62L^high^ MPPs (**Figure S11a-b**), a result also supported by a report in the literature that MEL-14 inhibits CD62L function on leukocytes^52^. To overcome this limitation, we lentivirally barcoded total MPPs and transplanted them into irradiated recipient mice (**Figure 5a**).Barcodes present in the CD62L^neg^ and CD62L^high^ MPPs, as well as the nucleated erythroid, B cells and myeloid cells were analysed 3 week post-transplantation – the timepoint when myeloid production from MPPs peaks post-transplantation^47^ (**Figure 5a-b**). We obtained 172 barcodes from all the samples that passed QC and filtering with high consistency of sequencing read counts between PCR technical duplicates and very little sharing of barcodes between mice (**Figure S12b-d**). When comparing the distribution of barcodes between CD62L^neg^ and CD62L^high^ MPPs, we found 124 barcodes that are shared between the two types of MPPs, suggesting that CD62L^low^ can give rise to CD62L^high^ MPPs and vice-versa or that cells may transition directly between compartments without undergoing cell division (**Figure 5b**). We also observed barcodes present in only one of the two MPP subsets (**Figure 5b**). Focusing on the differentiation outcome of the barcodes that had more than 95% of its reads in either the CD62L^low^ (CD62L^low enriched^; 30 barcodes) or the CD62L^pos^ (CD62L^high enriched^; 18 barcodes) MPPs, we found that CD62L^high enriched^ barcodes produce more myeloid cells and less B cells compared to CD62L^low enriched^ barcodes, while erythroid production was similar (**Figure 5c**).When comparing the lineage bias score of CD62L^neg enriched/high enriched^ barcodes, we found that CD62L^high enriched^ MPPs had a significantly higher myeloid bias than CD62L^low^ MPPs and significantly reduced B-cell bias and similar erythroid bias (**Figure 5d**). These results were corroborated by unsupervised clustering of the data where CD62L^high enriched^ barcodes cluster most closely with the myeloid lineage than the CD62L^low enriched^ indicating that CD62L^high^ HSPCs produce more myeloid cells (**Figure 5e**). Importantly, the total number of unique barcodes detected was similar for both the CD62L^high^ and the CD62L^neg^ subsets (**Figure S13a**). This confirmed that the reduced number of CD62L^neg^ barcodes in myeloid cells could not be explained by sampling or sensitivity issues, or differences due to the relative engraftment rates of the subsets. Furthermore, similar patterns in myeloid bias for CD62L^pos^ MPPs was observed when we transplanted lentivirally barcoded CD150^+^ HSCs, and purified CD62L MPP subsets at 12 months post transplantation (**Figure S14**). Together, our cellular barcoding experiments show that CD62L^high^ MPPs play a key role in repopulating and maintaining myeloid cell numbers in transplantation hematopoiesis. In summary, our infection and transplantation experiments show that metabolically-primed multipotent progenitors play a key role in fuelling emergency myelopoiesis.

**Figure 5:**
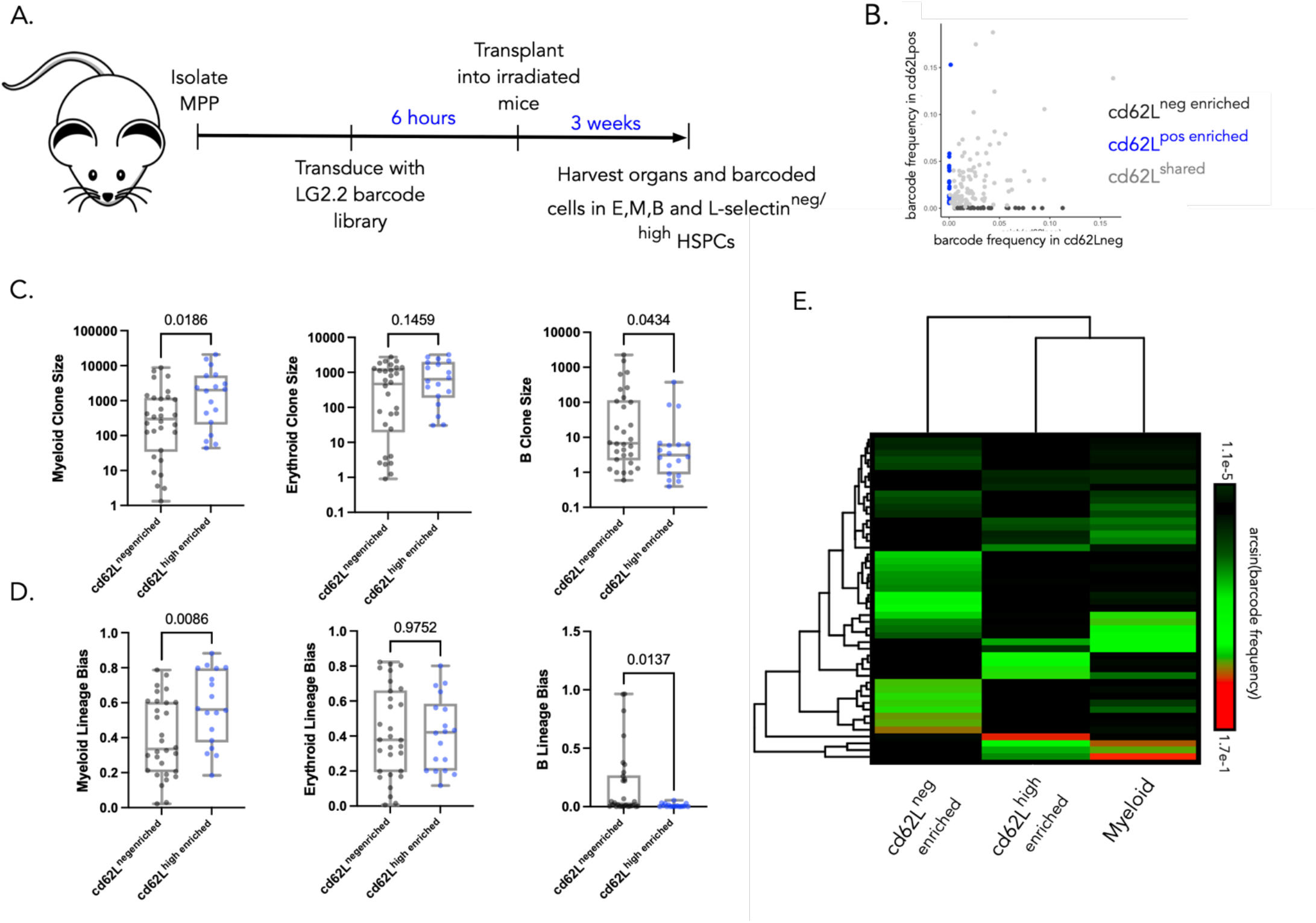
The CD62L^hi^ MPP compartment reconstitutes the myeloid compartment following bone marrow transplantation. (A) MPPs were purified from male donor B6 WT mice by FACs and infected with the LG2.2 lentiviral barcoding library for 6 hours. Transduced cells were transplanted into 4 sublethally irradiated (6Gy) control recipient mice (male littermate controls) and 3 weeks later CD62L^neg/hi^ MPPs, CD19^+^ B cells, CD44^+^ Ter119^+^ erythrocytes, and CD11b^+^ myeloid cells, as well as CD62L^high^ or ^neg^ MPPs from the bone marrow were sorted and processed for targeted sequencing of lentiviral lineage barcodes. (B) read abundance of barcode in the CD62L^neg enriched^ and the CD62L^high enriched^ HSPC (LSK) fraction, each dot is a barcode and the axis are transformed using the hyperbolic arcsin function. Barcodes that had more than 95% of its reads in either the CD62L^low^ or the CD62L^pos^ MPPs were classified as CD62L^shared^ (light grey; 124 barcodes), CD62L^neg enriched^ (dark grey; 30 barcodes) and CD62L^high enriched^ (blue; 18 barcodes) MPPs. (C) Number of cells of a given lineage (the myeloid, erythroid and B-cell lineages) produced per barcode, is clone size, for the barcode categories CD62L^neg enriched^ (grey) and CD62L^high enriched^ (blue) MPP subsets. Each point represents a distinct barcode. Statistical comparisons were made using a Mann-Whitney test. (D) lineage bias value for the myeloid, erythroid and B-cell lineages for barcodes in CD62L^neg enriched^ (grey) and CD62L^high enriched^ (blue) MPP subsets. Lineage bias represents the relative frequency of each barcode across the 3 mature cell lineage. Each point represents a distinct barcode. Statistical comparisons were made using a Mann-Whitney test. (E) Unsupervised clustering (using the Euclidean distance) and heatmap visualisation showing hyperbolic arcsin transformed barcode abundances for CD62L^neg enriched^ and CD62L^high enriched^ barcodes. Every row is a barcodes and every column is a cell type. All Boxplots represent the median and interquartile range with whiskers extending to the minimum and maximum values. N= 4 mice.

## Discussion

A key goal for the field of stem cell biology is to identify the molecular signals that induce stem cells to selectively differentiate into a cell type of interest *in vivo*. Previous work has highlighted the critical role of transcription and growth factors in regulating lineage commitment, but the role of metabolism is less clear in this context. In this study we developed MetaFate to trace the metabolic state and developmental fate of single HSPCs *in vivo*. Using this innovative approach, we characterise the metabolic cues that instruct early myeloid development, showing that the pentose phosphate pathway plays an active role in this process and can be manipulated to alter the rate of immune cell regeneration.

To add a functional dimension to metabolic studies of rare cell types *in vivo* we developed MetaFate, a lineage tracing approach to perform state-fate mapping focused on metabolically-associated gene modules. Within a single mouse, there are an estimated 1.4 x 10^5^ MPPs, representing just 0.03% of total bone marrow cellularity^2^. Consequently, cell yields falling far below the sensitivity limits of many metabolomics methods, making it technically challenging to study the metabolic properties of these rare subsets. Furthermore, existing methods are destructive – making it difficult to link the metabolic and functional properties of heterogeneous cell populations. Because of these technical challenges, much of our understanding of stem cell metabolism comes from population based approaches, limiting our understanding of how metabolism regulates lineage commitment specificity. MetaFate can also interface with other metabolomics methods – identifying functionally resolved sets of enzymes and transporters that can be verified at the protein level using spatially resolved metabolomics methods such as high dimensional mass cytometry^23^ or by *in situ* dehydrogenase assays^26^. We anticipate that combining emerging metabolomics technologies such as SCENITH and SPICE-Met with lineage tracing tools like MetaFate will yield significant insights into how cellular metabolism regulates the function of rare cell types in both health, ageing and disease. Given the diverse array of lineage tracing and metabolomics technologies that are emerging, our strategy can be readily adapted to other stem cell and developmental systems, including human tissues using human-compatible retrospective lineage tracing methods.

Using MetaFate, we have identified an expression program of enzymes and transporters that confers differences in myeloid lineage potential within a subset of MPPs. Leveraging the ability of the DRAG barcoding system to detect barcodes at both the RNA or the DNA level we were able to measure barcode abundances in both HSPCs and the much larger mature myeloid and nucleated erythroid progenitor compartments. This experimental design enabled us to trace lineage commitment over much longer developmental trajectories compared to studies that measure barcodes only in progenitors^27,37^. Using only genes encoding metabolic enzymes and transporters, our signature had a higher correlation with myeloid bias than the existing MPP3 signature^34,53^, prompting us to develop a novel purification strategy using the surface marker CD62L. Through *in situ* and lentiviral barcoding experiments, we show that CD62L enriches for myeloid bias in MPPs. This is consistent with reports showing that CD62L enriches for MPPs, rather than HSCs within the LSK compartment^54^ and that CD62L enriches for myeloid potential in CMPs^55^. In humans, the CD62L gene *SELL* has been associated with abnormal myeloid cell counts^56,57^, suggesting that CD62L may have implications in the regulation of hematopoiesis in humans as well as mice. Metabolically, CD62L^high^ MPPs have higher rates of ATP turnover and protein synthesis compared to CD62L^neg^ MPPs. CD62L^high^ MPPs also have a higher dependence on glucose metabolism via OXPHOS to meet their energetic requirements compared to other MPP subsets which had a higher reliance on glycolysis and on fatty/amino acid oxidation.

Importantly, manipulating metabolic processes in progenitors alters the rate of immune cell production, with overexpression of G6PD altering the dynamics of B-cell producing MPPs, resulting in a net skewing towards the erythromyeloid lineages. This result is consistent with reports that pharmacological inhibition of the pentose phosphate pathway blocks erythropoiesis *in vitro^8^* and that the pathway regulates the function of dendritic cells^58^ and macrophages^59,60^. This work bridges understanding between the fields of metabolism, stem cell biology and immunology, highlighting the pentose phosphate pathway as a regulator of immune cell production. While much focused has been placed on the roles of glycolysis and oxidative phosphorylation as key modulators of stem cell metabolism^14,15^, further work is required to assess whether the pentose phosphate pathway regulate can regulate stem cell function in other systems.

To understand why metabolic priming of multipotent progenitors may be functionally important we assessed the role of the CD62L^high^ MPP compartment in 2 different emergency myelopoiesis models: infection and transplantation. In both models, myelopoiesis was fuelled by CD62L^high^ MPPs, consistent with the idea that metabolic priming of multipotent progenitors toward the myeloid lineage facilitates the production of innate immune cells in response to injury. Our work therefore shows for the first time that MPPs are metabolically heterogeneous and that a subset of metabolically primed MPPs contribute to innate immunity in emergency settings.

In our study, we use transcriptomic changes that occur during fate decisions to infer metabolic differences that we validate using metabolic pathway activity measures. Differences in metabolite levels or others changes not captured by transcriptomic analysis may also regulate fate decisions. In particular, changes in metabolites acting as substrates for chromatin modifiers may precede transcriptomic changes and influence fate. Recent advances in single cell techniques to study the epigenome will help to address this limitation. In this context, metabolic differences could arise by cell extrinsic mechanisms such as differential location within the niche, or cell intrinsic mechanisms whereby the asymmetric distribution of metabolites/cell organelles following cell division could influence lineage potential. Further exploration of these topics is required to better understand the role of metabolism in shaping fate decisions.

Understanding the nutrients and metabolites that regulate hematopoiesis can inform the development of novel bone marrow organoid technologies to maintain and differentiate haematopoietic precursors *ex vivo*. Our data can also help to inform the development of nutrient/metabolite biomarker panels for stem cell function. Such tools can inform dietary interventions to promote HSPC function, particularly in prospective recipients of bone marrow transplants, a high-risk procedure that can lead to malnutrition and significant nutrient deficiencies^61,62^. Lastly, our results suggest that therapeutic interventions to alter the metabolism of HSPCs may not target all cells uniformly, given their underlying metabolic heterogeneity. Our approach may therefore be useful in studying whether this phenomenon occurs in other systems, such as cancer stem cells.

Collectively, we have identified the metabolic cues that guide the earliest stages of innate immune cell development, highlighting a key role for the pentose phosphate pathway. More broadly, our results suggest that manipulating lineage-specific metabolic cues can alter the cellular composition of the immune system *in vivo*.

## Materials and Methods

A detailed description of all materials and method is provided in the supplementary information

## Acknowledgements

We thank the Institute Curie flow cytometry, next-generation sequencing, animal, and UMR168 BMBC facility. We thank all the members of the Perié team for helpful discussion.

## Author Contributions

**J.C.** conceptualization, performed experiments, data curation, data analysis, methodology, writing, funding acquisition. **A.M.L.** data analysis, methodology, writing - review and editing. **I.R.** performed experiments **V.C.** data analysis, writing, review and editing. **C.C.** methodological development, performed experiments **S.T.B.** methodological development. **E.T.** methodological development. **E.R.** performed experiments. **F.T.** performed experiments. **Y.B.** performed experiments. **S.M.T.** performed experiments. **S.R.** performed experiments. **F.M.** performed experiments. **A.A.** provided expertise and reagents, writing, review and editing **C.L.** provided expertise and reagents, performed experiments, writing - review and editing. **P.B.** provided expertise and reagents, Writing - review and editing. **P.J.F.M.** provided expertise and reagents, writing - review and editing. **H.I.** Formal analysis, Writing - review and editing. **R.J.A.** provided expertise and reagents, writing - review and editing. **L.P.** conceptualization, data analysis, funding acquisition, methodology, supervision, writing.

## Ethics

All the experimental procedures were approved by the local ethics committee CEEA-IC (Comité d’Ethique en expérimentation animale de l’Institut Curie) under approval numbers DAP 2016 006, DAP 2021-010 and DAP 2021-013

## Data Availability

All datasets generated or reanalysed during this study are available at: https://github.com/TeamPerie/Cosgrove-et-al-2022

## Code Availability

All source code generated during this study is available at: https://github.com/TeamPerie/Cosgrove-et-al-2022

## References

1. Sender, R. & Milo, R. The distribution of cellular turnover in the human body. Nat. Med. 27, 45–48 (2021).

2. Cosgrove, J., Hustin, L. S. P., de Boer, R. J. & Perié, L. Hematopoiesis in numbers. Trends Immunol. S1471-4906(21)00211–8 (2021) doi:10.1016/j.it.2021.10.006.

3. Boettcher, S. & Manz, M. G. Regulation of Inflammation-and Infection-Driven Hematopoiesis. Trends Immunol. 38, 345–357 (2017).

4. Oburoglu, L., Romano, M., Taylor, N. & Kinet, S. Metabolic regulation of hematopoietic stem cell commitment and erythroid differentiation. Curr. Opin. Hematol. 23, 198–205 (2016).

5. Morganti, C., Cabezas-Wallscheid, N. & Ito, K. Metabolic Regulation of Hematopoietic Stem Cells. HemaSphere 6, e740 (2022).

6. Takubo, K. et al. Regulation of glycolysis by Pdk functions as a metabolic checkpoint for cell cycle quiescence in hematopoietic stem cells. Cell Stem Cell 12, 49–61 (2013).

7. Signer, R. A. J., Magee, J. A., Salic, A. & Morrison, S. J. Haematopoietic stem cells require a highly regulated protein synthesis rate. Nature 509, 49–54 (2014).

8. Oburoglu, L. et al. Glucose and glutamine metabolism regulate human hematopoietic stem cell lineage specification. Cell Stem Cell 15, 169–184 (2014).

9. Ito, K. et al. A PML–PPAR-δ pathway for fatty acid oxidation regulates hematopoietic stem cell maintenance. Nat. Med. 18, 1350–1358 (2012).

10. Ito, K. et al. Self-renewal of a purified Tie2+ hematopoietic stem cell population relies on mitochondrial clearance. Science 354, 1156–1160 (2016).

11. Cabezas-Wallscheid, N. et al. Vitamin A-Retinoic Acid Signaling Regulates Hematopoietic Stem Cell Dormancy. Cell 169, 807–823.e19 (2017).

12. Agathocleous, M. et al. Ascorbate regulates haematopoietic stem cell function and leukaemogenesis. Nature 549, 476–481 (2017).

13. Qi, L. et al. Aspartate availability limits hematopoietic stem cell function during hematopoietic regeneration. Cell Stem Cell 28, 1982–1999.e8 (2021).

14. Ly, C. H., Lynch, G. S. & Ryall, J. G. A Metabolic Roadmap for Somatic Stem Cell Fate. Cell Metab. 31, 1052–1067 (2020).

15. Shapira, S. N. & Christofk, H. R. Metabolic Regulation of Tissue Stem Cells. Trends Cell Biol. 30, 566–576 (2020).

16. Busch, K. et al. Fundamental properties of unperturbed haematopoiesis from stem cells in vivo. Nature 518, 542–546 (2015).

17. Naik, S. H. et al. Diverse and heritable lineage imprinting of early haematopoietic progenitors. Nature 496, 229–232 (2013).

18. Perié, L., Duffy, K. R., Kok, L., de Boer, R. J. & Schumacher, T. N. The Branching Point in Erythro-Myeloid Differentiation. Cell 163, 1655–1662 (2015).

19. Shamir, M., Bar-On, Y., Phillips, R. & Milo, R. SnapShot: Timescales in Cell Biology. Cell 164, 1302–1302.e1 (2016).

20. DeVilbiss, A. W. et al. Metabolomic profiling of rare cell populations isolated by flow cytometry from tissues. eLife 10, e61980 (2021).

21. Sun, J. et al. Clonal dynamics of native haematopoiesis. Nature 514, 322–327 (2014).

22. Pei, W. et al. Polylox barcoding reveals haematopoietic stem cell fates realized in vivo. Nature 548, 456–460 (2017).

23. Hartmann, F. J. et al. Single-cell metabolic profiling of human cytotoxic T cells. Nat. Biotechnol. (2020) doi:10.1038/s41587-020-0651-8.

24. Argüello, R. J. et al. SCENITH: A Flow Cytometry-Based Method to Functionally Profile Energy Metabolism with Single-Cell Resolution. Cell Metab. 32, 1063–1075.e7 (2020).

25. Russo, E. et al. SPICE-Met: profiling and imaging energy metabolism at the single-cell level using a fluorescent reporter mouse. EMBO J. e111528 (2022) doi:10.15252/embj.2022111528.

26. Miller, A. et al. Exploring Metabolic Configurations of Single Cells within Complex Tissue Microenvironments. Cell Metab. 26, 788–800.e6 (2017).

27. Rodriguez-Fraticelli, A. E. et al. Single-cell lineage tracing unveils a role for TCF15 in haematopoiesis. Nature 583, 585–589 (2020).

28. Pei, W. et al. Resolving Fates and Single-Cell Transcriptomes of Hematopoietic Stem Cell Clones by PolyloxExpress Barcoding. Cell Stem Cell 27, 383–395.e8 (2020).

29. Cavicchioli, M. V., Santorsola, M., Balboni, N., Mercatelli, D. & Giorgi, F. M. Prediction of Metabolic Profiles from Transcriptomics Data in Human Cancer Cell Lines. Int. J. Mol. Sci. 23, 3867 (2022).

30. Sawai, C. M. et al. Hematopoietic Stem Cells Are the Major Source of Multilineage Hematopoiesis in Adult Animals. Immunity 45, 597–609 (2016).

31. Bowling, S. et al. An Engineered CRISPR-Cas9 Mouse Line for Simultaneous Readout of Lineage Histories and Gene Expression Profiles in Single Cells. Cell 181, 1410–1422.e27 (2020).

32. Wilson, N. K. et al. Combined Single-Cell Functional and Gene Expression Analysis Resolves Heterogeneity within Stem Cell Populations. Cell Stem Cell 16, 712–724 (2015).

33. Giladi, A. et al. Single-cell characterization of haematopoietic progenitors and their trajectories in homeostasis and perturbed haematopoiesis. Nat. Cell Biol. 20, 836–846 (2018).

34. Pietras, E. M. et al. Functionally Distinct Subsets of Lineage-Biased Multipotent Progenitors Control Blood Production in Normal and Regenerative Conditions. Cell Stem Cell 17, 35–46 (2015).

35. Tusi, B. K. et al. Population snapshots predict early haematopoietic and erythroid hierarchies. Nature 555, 54–60 (2018).

36. Dahlin, J. S. et al. A single-cell hematopoietic landscape resolves 8 lineage trajectories and defects in Kit mutant mice. Blood 131, e1–e11 (2018).

37. Weinreb, C., Rodriguez-Fraticelli, A., Camargo, F. D. & Klein, A. M. Lineage tracing on transcriptional landscapes links state to fate during differentiation. Science 367, (2020).

38. Wolf, F. A. et al. PAGA: graph abstraction reconciles clustering with trajectory inference through a topology preserving map of single cells. Genome Biol. 20, 59 (2019).

39. Paul, F. et al. Transcriptional Heterogeneity and Lineage Commitment in Myeloid Progenitors. Cell 163, 1663–1677 (2015).

40. Chen, C. et al. TSC–mTOR maintains quiescence and function of hematopoietic stem cells by repressing mitochondrial biogenesis and reactive oxygen species. J. Exp. Med. 205, 2397–2408 (2008).

41. Yu, W.-M. et al. Metabolic regulation by the mitochondrial phosphatase PTPMT1 is required for hematopoietic stem cell differentiation. Cell Stem Cell 12, 62–74 (2013).

42. Verny, L., Sella, N., Affeldt, S., Singh, P. P. & Isambert, H. Learning causal networks with latent variables from multivariate information in genomic data. PLOS Comput. Biol. 13, e1005662 (2017).

43. Choi, J. et al. Haemopedia RNA-seq: a database of gene expression during haematopoiesis in mice and humans. Nucleic Acids Res. 47, D780–D785 (2019).

44. Cabeli, V. et al. Learning clinical networks from medical records based on information estimates in mixed-type data. PLOS Comput. Biol. 16, e1007866 (2020).

45. Nóbrega-Pereira, S. et al. G6PD protects from oxidative damage and improves healthspan in mice. Nat. Commun. 7, 10894 (2016).

46. Eisele, A. S. et al. Erythropoietin directly remodels the clonal composition of murine hematopoietic multipotent progenitor cells. eLife 11, e66922 (2022).

47. Boyer, S. W. et al. Clonal and Quantitative In Vivo Assessment of Hematopoietic Stem Cell Differentiation Reveals Strong Erythroid Potential of Multipotent Cells. Stem Cell Rep. 12, 801–815 (2019).

48. Schultze, J. L., Mass, E. & Schlitzer, A. Emerging Principles in Myelopoiesis at Homeostasis and during Infection and Inflammation. Immunity 50, 288–301 (2019).

49. Boettcher, S. et al. Cutting edge: LPS-induced emergency myelopoiesis depends on TLR4-expressing nonhematopoietic cells. J. Immunol. Baltim. Md 1950 188, 5824–5828 (2012).

50. Haltalli, M. L. R. et al. Manipulating niche composition limits damage to haematopoietic stem cells during Plasmodium infection. Nat. Cell Biol. 22, 1399–1410 (2020).

51. Mitroulis, I., Kalafati, L., Hajishengallis, G. & Chavakis, T. Myelopoiesis in the Context of Innate Immunity. J. Innate Immun. 10, 365–372 (2018).

52. Pizcueta, P. & Luscinskas, F. W. Monoclonal antibody blockade of L-selectin inhibits mononuclear leukocyte recruitment to inflammatory sites in vivo. Am. J. Pathol. 145, 461–469 (1994).

53. Sommerkamp, P. et al. Mouse multipotent progenitor 5 cells are located at the interphase between hematopoietic stem and progenitor cells. Blood 137, 3218–3224 (2021).

54. Cho, S. & Spangrude, G. J. Enrichment of functionally distinct mouse hematopoietic progenitor cell populations using CD62L. J. Immunol. Baltim. Md 1950 187, 5203–5210 (2011).

55. Ito, Y., Nakahara, F., Kagoya, Y. & Kurokawa, M. CD62L expression level determines the cell fate of myeloid progenitors. Stem Cell Rep. 16, 2871–2886 (2021).

56. Orrù, V. et al. Complex genetic signatures in immune cells underlie autoimmunity and inform therapy. Nat. Genet. 52, 1036–1045 (2020).

57. Vuckovic, D. et al. The Polygenic and Monogenic Basis of Blood Traits and Diseases. Cell 182, 1214–1231.e11 (2020).

58. Everts, B. et al. TLR-driven early glycolytic reprogramming via the kinases TBK1-IKKε supports the anabolic demands of dendritic cell activation. Nat. Immunol. 15, 323–332 (2014).

59. Tannahill, G. M. et al. Succinate is an inflammatory signal that induces IL-1β through HIF-1α. Nature 496, 238–242 (2013).

60. Haschemi, A. et al. The sedoheptulose kinase CARKL directs macrophage polarization through control of glucose metabolism. Cell Metab. 15, 813–826 (2012).

61. The EBMT Handbook: Hematopoietic Stem Cell Transplantation and Cellular Therapies. (Springer Nature, 2019). doi:10.1007/978-3-030-02278-5.

62. Akbulut, G. & Yesildemir, O. Overview of nutritional approach in hematopoietic stem cell transplantation: COVID-19 update. World J. Stem Cells 13, 1530–1548 (2021).

