## supplementary information for "Metabolically Primed Multipotent Hematopoietic Progenitors Fuel Innate Immunity"

### Materials and Methods

**Mice:** All the experimental procedures were approved by the local ethics committee CEEA-IC (Comité d'Ethique en expérimentation animale de l'Institut Curie) under approval numbers DAP 2016 006, DAP 2021-010 and DAP 2021-013 and by the Institut Pasteur Safety Committee in accordance with French and European guidelines (CETEA 190148). Within each experiment mice were sex and aged matched and littermates were used to generate control cohorts. SCENITH and LPS challenge experiments were performed in C57BL6/J mice. Transplantation studies were performed in either CD45.1 C57BL6N / CD45.2 C57B6N mice or in HG6PD(Tg+) or HG6PD(Tg-) littermate controls. G6PD-Tg mice and littermate controls were generated the Spanish National Cancer Research Center (CNIO) at the Transgenic Mice core facility and were provided by Pablo Jose Fernández-Marcos. PercevalHR Vav-iCre mice were generated by the Centre d'Ingénierie Génétique Murine at Institut Pasteur and performed in accordance with the European Community guidelines (2010/63/UE) and were provided by P. Bousso. For DRAG barcoding experiments DRAG1 mice were crossed with B6.Cg-Tg(CAG-cre/Esr1*)5Amc/J (CAGGCre-ER$^{TM}$) to obtain heterozygous mice.

**Cell isolation:** BM cells were obtained from wild-type C57BL/6 of 8-16 weeks of age by bone flushing of femur tibia and iliac crest. Bone marrow cells were MACS enriched for cKit+ cells using CD117 MicroBeads Ultrapure (Miltenyi Biotec cat #130-091-224) according to manufacturer's protocol. Cells were kept in RPMI + 10% fetal calf serum + 1% pen-strep or in PBS + 10% fetal-calf serum + 1% pen-strep.

**Flow Cytometry:** Cell suspensions were incubated with antibody master mixes for 30 minutes on ice protected from ambient light. Cells were then washed in RPMI + 10% FCS + 1% pen-step for 2 x 5 minutes for 1700rpm at 4C. For TMRE staining of mitochondrial membrane potential cell suspensions were processed for flow cytometry surface staining and were then incubated in 200ul of 100nM TMRE solution. A full list of the antibodies used in this study is provided in table S1. Samples were processed on a Ze5 (bio-rad), or Cytoflex LX (Beckman Coulter) plate-reader cytometers. Data analysis was performed using FlowJo v10.2 software (TreeStar), R v4.2.0, and Prism v9.

**Fluorescence activated cell sorting:** FACS was performed at the flow cytometry facility of Institute Curie on a FACSAria$^{TM}$ (BD Biosciences) or sh800 (Sony). Cells were sorted using a 70 µm nozzle at precision 0/16/0 and high efficiency in epperdendorf tubes.

**SCENITH:** cKit enriched murine bone marrow cells were seeded in 96 well plates for studying blood cells metabolism at 2 × $10^6$ cells/mL. Wells were treated during 30-60 minutes with Control, Oligomycin (Oligo, final concentration 1 µM), or the translation initiation inhibitor Harringtonine. 2-Deoxy-D-Glucose (DG, final concentration 100mM), was not used in our final analysis as treatment with this molecule for long time-periods (> 30 minutes) along with fixing and permeabilization of the cells led to shedding of CD62L from the membrane of MPPs. This effect was not observed in SPICE-Met processed cells, which are exposed to the same concentration of 2-DG, but are measured immediately after staining in live cells, without fixation and permeabilization steps. Puromycin (final concentration 10 µg/mL) is added at the same time as the metabolic inhibitor treatment. After puromycin treatment, cells were washed in cold PBS and stained with a combination of Fc receptors blockade and fluorescent cell viability marker (fixable live/dead nearIR, ThermoFisher Scientific, cat # L10119), then primary conjugated antibodies against surface markers (**table S1**) during 25 min at 4°C in PBS 1X 5% FCS, 2mM EDTA (FACS wash buffer). After washing, cells were fixed and permeabilized using FOXP3 fixation and permeabilization buffer (Thermofisher eBioscience) following manufacturer instructions. Intracellular staining of puro using fluorescently labeled anti-Puro monoclonal antibody with Alexa Fluor 488 was performed by incubating cells during 1 h at 4°C diluted in permeabilization buffer. Mitochondrial dependencies were calculated as $((\text{Puro-MFI}_{DMSO+H2O} - \text{Puro-MFI}_{oligo}) / (\text{Puro-MFI}_{DMSO+H2O} - \text{Puro-MFI}_{H}))$ x100.

**SPICE-Met:** cKit enriched cells were purified from the femur, tibia and iliac bones of vav-iCre Perceval$^{fl/fl}$ mice. Following surface staining with antibodies (**table S1**), cells expressing PercevalHR were treated with either 1mM oligomycin, 100 mM 2-deoxy-D-glucose (2-DG), the combination of the two or DMSO as control and kept at 37°C for 30 minutes and then analyzed by flow cytometry. Samples were then recorded for one minute on a Cytoflex LX (Beckman Coulter). ATP:ADP ratios in individual cells were calculated from two fluorescence signals. In brief, excitation with a violet laser (405nm) and signal detection with a 525 band-pass filter was used to measure ADP contribution while ATP levels were estimated by excitation with a blue laser (488nm) and signal detection with a 525 band-pass filter. Data analysis was performed using FlowJo. OXPHOS and glucose contribution were calculated as $((R_{DMSO+H2O} - R_{oligo}) / (R_{DMSO+H2O} - R_{oligo+2\text{-}DG}))$ x100 and $((R_{DMSO+H2O} - R_{2\text{-}DG}) / (R_{DMSO+H2O} - R_{oligo+2\text{-}DG}))$ x100, respectively where R represents the ratio of fluorescent intensity measurements for ATP and ADP .

### <u>Lentiviral Barcoding</u>

**Lentiviral Barcoding Transduction**: The barcode library LG2.2 was used as in Eisele *et al* (2022) [1]. In brief, lentiviruses were produced by transfecting the barcode plasmids and p8.9-QV and pVSVG into HEK293T cells in DMEM-Glutamax supplemented with 10% FCS (Gibco), 1% MEM NEAA, and 1% sodium pyruvate using Polyethyleneimine. Supernatant was 0,45 um filtered, concentrated by 1h30 ultracentrifugation at 31 000g and frozen at -80°C. For isolation and labelling of cells with the LG2.2 lentiviral barcoding library bone marrow cells were isolated from femur, tibia and iliac bones by flushing using a 21G needle (Terumo), and C-Kit$^{+}$ cells enriched using anti-CD117 magnetic beads (Miltenyi) on the MACS column system (Miltenyi). Cells were stained with surface antibodies (**table S1**) and sorted at the flow cytometry facility of Institute Curie on a FACSAria$^{TM}$ (BD Biosciences) or sh800 (Sony). Cells were sorted using a 70 μm nozzle at precision 0/16/0 and high efficiency in epperdendorf tubes. MPPs were transduced with the lentiviral barcode library in StemSpanMedium SFEM (STEMCELL Technologies) supplemented with 50 ng/ml mSCF (STEMCELL Technologies) through 1,5 h of centrifugation at 300 g followed by 4,5 h incubation at 37°C in order to obtain 10% barcoded cells. After the incubation 15-18,000 cells were injected in the tail vein of 6 Gy sub-lethally irradiated recipient mice.

**Lentiviral Barcode amplification and sequencing:** Bones (femurs, tibias and ilia) were isolated for barcode analysis from recipient mice. Bone marrow cells were extracted by flushing of the bones using a 21G needle (Terumo) and enriched using anti-CD117 magnetic beads (Miltenyi) on the MACS column system (Miltenyi). The ckit$^{-}$ fraction was further separated into a Ter119$^{+}$ and Ter119$^{-}$ fractions using anti-Ter119 coated magnetic beads (Miltenyi). Cells were stained with fluorescently conjugated antibodies (table S1) and after sorting mature cells were lysed in 40 µl Viagen Direct PCR Lysis Reagent (cell) (Euromedex) supplemented with 0,5 mg/ml Proteinase K Solution RNA grade (Invitrogen) in a thermic cycler: (55°C for 120 min, 85°C for 30 min, 95°C for 5 min, indefinite at 4°C). Samples were then split into two replicates, and a three-step nested PCR was performed to, in a first step amplify barcodes (primers top-LIB (5'TGCTGCCGTCAACTAGAACA-3') and bot-LIB (5'GATCTCGAATCAGGCGCTTA-3')), in a second step add unique 4 bp plate indices (forward 5'ACACTCTTTCCCTACACGACGCTCTTCCGATCTNNNNCTAGAACACTCGAGATCAG3' and reverse 5'GTGACTGGAGTTCAGACGTGTGCTCTTCCGATCGATCTCGAATCAGGCGCTTA3'), and in a third step add P5 and P7 flow cell attachment sequences and one of 96 sample indices of 7 bp P5 5'AATGATACGGCGACCACCGAGATCTACACTCTTTCCCTACACGACGCTCTTCCGATCT3' and P7 5'CAAGCAGAAGACGGCATACGAGANNNNNNNGTGACTGGAGTTCAGACGTGCTCTTCCGATC3') (PCR program: hot start 5 min 95°C, 15 s at 95°C; 30 s at 57.2°C; 30 s at 72°C, 5 min 72°C, 30 (PCR1-2) or 15 cycles (PCR 3)). Both index sequences (sample and plate) were designed based on[2] such that sequences differed by at least 2 bases, and homopolymers or more than 2 bp, hairpins and complementary regions with the rest of the primer sequence were absent. To avoid lack of diversity at the beginning of the reads during sequencing, at least 4 different plate indices were used for each sequencing run. Primers were ordered desalted, and high-performance liquid chromatography (HPLC) purified. During lysis and each PCR, a mock control was added. The DNA amplification by the three PCRs was monitored by the run on a large 2% Agarose gel. Samples were pooled in order to guarantee a sequencing depth of 50 reads/cell. Five µl of the products of PCR3 for each sample and replicate were pooled, purified using the Agencourt AMPure XP system (Beckman Coulter), analyzed on a Bioanalyzer, and diluted to a concentration of 5 nM. These pools were sequenced on a HiSeq system (Illumina) (SR-65bp) at the sequencing facility of Institute Curie (10% of Phix Illumina phage genome library was added to generate a more diverse set of clusters).

**Lentiviral Barcode Analysis:**

Sequencing results were analyzed using R-4.2.0, Microsoft Excel (v16.16, MAC edition), and GraphPad Prism version 9.0.

**Data demultiplexing:** Reads were first filtered for perfect match to the input index- and common-sequences using XCALIBR (https://github.com/NKI-GCF/xcalibr) and filtered against a barcode reference list.

**Data QC:** The consistency of technical replicates for each sample were then assessed using a Pearsons Correlation but no filtering was applied based on this metric. Barcodes that were not present in both technical replicates were filtered from the data with 3606/3969 WT barcodes and 3461/3942 G6PD-Tg barcodes retained at this step. We then assessed the prevalence of repeat-use barcodes in our dataset. In this context barcodes that occur in more than one mouse per transduction batch are likely to be the result of more than 1 cell being labelled with the same barcode. To estimate repeat-use frequency in our datasets we first had to ascertain which repeat use barcodes were the result of repeat labelling versus those that arose due to sequencing errors. To do this, we ranked repeat use barcodes based on how many sequencing reads mapped to each. Barcodes below the upper 95$^{th}$ percentile of read abundance are considered as sequencing noise and set to zero in each respective sample. Barcodes which are in the upper 95$^{th}$ percentile of read abundance and only found in one mice after noise correction are retained for further analysis. Following this filtering step 819 unique WT barcodes and 805 unique G6PD-Tg barcodes were retained for analysis.

**Data normalisation:** After filtering, we normalised barcode counts within each sample. In a barcode count matrix where rows are barcodes and columns are samples, we normalize the data so that each column sums to 1. More precisely, Let $R_{bc}$ represent the number of reads for barcode *B* in cell type *C,* and let $P_{bc}$ represent the proportional read abundance per barcode per cell type:

$$P_{bc} = \frac{R_{bc}}{\sum_b R_{bc}}$$

When calculating clone sizes for each barcode it is relevant to know the number of cells produced and so we scale normalized sequencing reads by the number of cells in each sample, giving the cell-scaled barcode frequency $C_{bc}$. During data acquisition, we do not always measure the entire tissue sample. In this setting the number of cells sorted by FACS is corrected by accounting for the fraction of the total sample that was measured. As a concrete example if a bone marrow cell suspension was placed in 5ml of medium but only 4ml of this medium was acquired during cell sorting we scale the number of cells sorted by the inverse of the proportion of the sample that was measured. (1/0.8 = 1.25 in this example)

$$C_{bc} = P_{bc}\, x\, (number\ of\ cells\ sorted * \frac{1}{fraction\ of\ total\ sample\ measured})$$

**Calculation of barcode diversity and clone sizes:**

Barcode diversity was calculated as the total number of unique barcodes found in a given sample, and where relevant the number of unique barcodes found in 2 developmentally related samples, for example CD62Lhi MPPs and myeloid cells. Clone sizes were calculated from the cell-scaled barcode frequencies ($C_{bc}$) of a given barcode in a given sample.

**Calculation of lineage bias score:** To classify barcodes by their lineage bias, an additional normalization step per barcode is applied in each individual, thereby enabling categorization of each barcode into classes of biased output towards the analyzed cell types. Let lineage-bias$_{bc}$ represent the relative representation of barcode *B* in cell type *C,* and let $P_{bc}$ represent the proportional read abundance per barcode per cell type

$$lineage\ bias_{bc} = \frac{P_{bc}}{\sum_c P_{bc}}$$

**Calculation of production bias:** In instances where the number of cells produced per progenitor is relevant to the assessment of bias, we perform the same calculation as for lineage bias but use cell-scaled barcode frequencies $C_{bc}$ as opposed to the proportional read abundance per barcode per cell type.

$$Production\ bias_{bc} = \frac{C_{bc}}{\sum_c C_{bc}}$$

### MetaFate

**Induction of Barcode Recombination**

8-11 weeks old male mice received 7 mg Tamoxifen/ 40 gr bodyweight each day for 3 consecutive days by intraperitoneal injection. Tamoxifen (T5648-1G, Sigma) was dissolved 10% EtOH and 90% sunflower oil.

**Cell isolation and sorting**

At sacrifice, BM was harvested from femurs, tibias and ilia and enriched using anti-CD117 magnetic beads (Miltenyi). The c-kit$^+$ fraction was stained with antibodies against CD117 (c-kit APC, clone 2B8, Biolegend), Sca-1 (Pacific Blue, clone D7, eBioscience). The c-kit$^-$ fraction was stained with antibodies against CD11b (PercPCy5.5 or Pacific Blue, clone M1/70, eBioscience), CD19 (APC-Cy7, clone 1D3, BD Pharmingen). FACS was performed at the flow cytometry facility of Institute Curie on a FACSAria$^{TM}$ (BD Biosciences) or sh800 (Sony). Data analysis was performed using FlowJo$^{TM}$ v.10 (TreeStar). Cells were sorted using a 70 μm nozzle at precision 0/16/0 and high efficiency.

**10X Genomics 3’ Library preparation for single-cell transcriptomics analysis:** DRAG barcoded LSK (ckit$^+$ sca1$^+$ gfp$^+$) cells were sorted from ckit-enriched bone marrow fraction. Samples were then processed using the 10X genomics Chromium Single Cell 3’ v3 kit. Specifically, 1,000-16,000 cells were loaded for each experiment for a targeted recovery of 500-10,000 cells. cDNA amplification was performed with 11-13 PCR cycles depending on the targeted cell recovery, as per the manufacturers recommendations. Sequencing was performed on a NovaSeq (illumina) on paired end PE28-8-91 mode. Raw sequencing reads were processed using Cellranger.

**Targeted RNA-Barcode Recovery from 3’ scRNA-seq libraries:** 10ng of our cDNA libraries underwent a cDNA pre-amplification using the 10X 3’ amplification mix, partial read1 F primer (1uM), PreAmp + read2 reverse primer (1uM) and $H_20$ and were incubated at 98C for 3 minutes followed by 15 cycles of 15 seconds at 98C, 20 seconds at 63C and 60 seconds at 72C, samples were then incubated for 60 seconds at 72C. Samples then underwent SPRI size selection at a ratio of 0.6X before undergoing a sample index PCR. 10ul of our preamplification product was placed in a master mix of 10X 3’ amplification mix along with P5- read 1 forward primer (1uM) and P7 reverse primer (1uM). Each sample then underwent PCR amplification with incubation at 98C for 45 seconds, followed by 8-16 cycles (according to table 1) of 98C for 20 seconds, 63C for 30 seconds, and 72C for 20 seconds. samples then underwent a 60 second incubation at 72C. Samples then underwent an additional SPRI cleanup at 0.8X. Sequencing was performed on a MiSeq (Illumina) on paired-end 300bp mode.

| cDNA Input | Total cycles (10x protocol) |
|---|---|
| 0,25-25ng | 16-18 |
| 25-150ng | 14-16 |
| 150-500ng | 12-14 |
| 500-1000ng | 10-12 |
| 1000-1500ng | 8-10 |
| >1500ng | 7 |

Table 1. PCR amplification cycles for targeted barcode recovery at the RNA level

**Targeted DNA-Barcode using PCR and deep sequencing on bulk mature cells.**

**Lysis:** Sorted cells were lysed 40 ul DirectPCR Lysis Reagent (Cell) from Viagen Biotech with 0.4 mg Prot K, and incubated for 1 h at 55$^{o}$C, followed by 30 min at 85$^{o}$C heat inactivation and 5 min at 94$^{o}$C. Samples were stored at -20$^{o}$C. **Preamp PCR:** Samples were resuspended in 200 µl PCR mix (20 µl 5x phusion HF buffer (NEB); 1 µl Phusion DNA Polymerase (NEB); 2 µl 10 mM dNTP's; 0.5 µl 100 uM preamp forw. oligo; 0.5 µl 100 uM preamp rev. oligo; 76 µl PCR grade water) and split in two replicates. PCR program: 2 min. at 98ºC; N* cycles of 10 sec at 98ºC, 20 sec at 60ºC, 25 sec at 72ºC; 5 min. at 72ºC; hold at 4ºC (* N is adjusted to number of barcoded cells in the sample, so at the moment of tagging every sample has the same number of molecules (See table S2). **Tagging PCR:** (adjusted from Kinde et al, PNAS, 2011)**:** 2 µl preamp PCR product was mixed with 48 µl tagging PCR mix (10 µl 5x phusion HF buffer (NEB); 1 µl Phusion Hot Start II DNA Polymerase (2U/ µl) (NEB); 1 µl 10 mM dNTP's; 0.25 µl 1 µM M1 tag forward oligo; 35.75 µl PCR grade water). PCR program: 1 min. at 98ºC; 2 cycles of 10 sec at 98ºC, 2 min. at 57ºC, 1 min. at 72ºC; 20ºC forever). To digest remaining M1 tag forward oligo: 3 µl 20U/ µl Exonuclease I (NEB) was added and incubated for 1 h at 37ºC, denaturing ExoI was done for 5 min at 98ºC, and the sample cooled down to 20ºC. 0.5 µl 100 µM Illumina forward seq oligo and M1rev oligo (both had 2 Phosphorothioate bonds at the 3'end to avoid breakdown from residual ExoI activity) were added, followed by PCR program: 1 min. at 98ºC; 30 cycles of 10 sec at 98ºC, 20 sec at 67ºC, 25 sec at 72ºC; 5 min. at 72ºC; 4ºC forever. **Sample index PCR:** 2 µl tagging PCR product was mixed with 18 µl PCR mix (4 µl 5x phusion HF buffer (NEB); 0.4 µl Phusion DNA Polymerase (NEB); 0.4 µl 10 mM dNTP's; 0.1 µl 100 uM P5 forw. oligo; 4 µl 2.5 uM P7 index rev. oligo; 9.1 µl PCR grade water). PCR program: 30 sec at 98ºC; 15 cycles of 10 sec at 98ºC, 20 sec at 67ºC, 25 sec at 72ºC; 5 min. at 72ºC; 4ºC forever). **NGS sequencing:** 5 µl from each index PCR was taken and pooled before being cleaned using a SPRI selection (ratio 1:1) The pooled samples were deep-sequenced on a MiSeq System (Illumina) in SR150bp run mode

**RNA DRAG Barcode Preprocessing and Filtering:** Cell Ranger was used to process the reads from the targeted sequencing, aligning them to the mouse genome and adding 10x cell and unique molecular identifier (UMI) barcodes. Unmapped reads were then extracted from the original 10x and targeted bams using 'samtools view -f 4' and concatenated. An exact grep match to 52 base pairs at the 5 prime (V) (corresponding to the targeted amplification primer) end of the DRAG lineage barcode plus 10 base pairs at the 3 prime (J) end, allowing 10-40 variables bases in between, was used to extract reads potentially containing a lineage barcode. This was designed such that the start and end of the variable bases matches the barcodes extracted from the DNA. These reads were further filtered to keep only those with a 10x cell barcode and UMI assigned, and then to keep only reads in cells (as defined by Cell Ranger).

To assign one VDJ barcode to each 10x cell barcode, we iterated through each 10x cell barcode, extracting all relevant reads. Then, UMIs were filtered to keep only those with 3 or more reads and one dominant VDJ barcode (defined as >=0.45 of total reads of that DRAG barcode for a given cell). The dominant barcode for each UMI was extracted, and finally we assign one VDJ barcode to a 10x cell if we have good agreement across UMIs, defined as >=0.75 of UMIs for that cell mapping to one VDJ sequence. If there is only one UMI retained, we further ensure that the VDJ barcode for this UMI is the dominant barcode across all the reads for that cell and has >=0.45 of reads.

As with VDJ barcodes extracted from DNA, we perform further checks and filters to ensure the barcodes are of good quality and unique. First we check that the barcodes conform to the expected VDJ structure, using an algorithm (Urbanus and Cosgrove, manuscript attached in supplementary information) to compare barcodes to the original VDJ template and identify which nucleotides were deleted due to exonuclease activity and which ones were inserted due to Tdt activity. This enabled us to recognize barcodes containing residual error, and also to quantify barcode creation patterns (i.e. the number of deleted and inserted nucleotides at the junctions between V, D and J segments per barcode). We used this approach to filter barcodes that did not have the expected barcode recombination pattern (this only removes ~1% of barcodes). We then use the iGoR algorithm to compute a generation probability for each barcode (Urbanus and Cosgrove, manuscript attached in supplementary information).

**DNA Barcode Preprocessing and Filtering.**

Each recombined sequence includes nucleotide additions and deletions (referred to as the 'DRAG barcode') and constant parts that flank both sides of this barcode. Moreover, each barcode was associated with a random unique molecular identifier (UMI) of 12bp during the tagging PCR step.

**DNA DRAG barcode Preprocessing.** We use the pipeline described below to demultiplex fastq files and identify the reads that match a potential recombination of the DRAG construct. First the bcl2fastq (Illumina) program is used to demultiplex the fastq files based on the i7 index sequence. Only records that match the i7 index perfectly are considered for the next step. In the constant part of the V and the J, the reads tend to be error-prone and a consensus sequence with Ns is manually created. The Xcalibr program (https://github.com/NKI-GCF/xcalibr) is then used to extract counts for all combinations of the 16bp UMI and the recombined barcodes only for the reads that contain the constant sequence of the V at the expected coordinates. After this, the J constant part (tagcaagctcgagagtagacctactggaatcagaccgccaccatggtgagc) is aligned to the barcode part using the NBCI blast2 program[3]. When a suitable match is found, the barcode is trimmed at the start coordinate of the match, resulting in the final matrix.

**DNA DRAG Barcode Processing.** We used the steps described below to identify barcode sequences and remove PCR and deep-sequencing errors. First, we removed any barcode and associated UMI containing one or multiple 'N' values (within either the barcode, constant flanking parts or UMI). Second, barcodes that did not have an exact match to the expected constant parts of the V and J that precede or follow the barcode were removed. Third, when multiple sequences were found associated with a single UMI, only the most frequently occurring barcode associated with that UMI was kept. As a fourth step, we summed up the read counts for all UMI associated with the set of remaining barcodes. To verify that the barcodes obtained match the expected structure of a VDJ recombination product, we developed an algorithm to compare barcodes to the original VDJ template and identify which nucleotides were deleted due to exonuclease activity and which ones were inserted due to Tdt activity. This enabled us to recognize barcodes containing residual error, and also to quantify barcode creation patterns (i.e. the number of deleted and inserted nucleotides at the junctions between V, D and J segments per barcode) (Urbanus and Cosgrove, manuscript attached in supplementary information).

**Single-Cell RNA-seq Analysis:** Raw sequencing reads were processed using Cellranger. To obtain a reads/cell/gene count table, reads were mapped to the mouse GRCm38.84 reference genome. During filtering, Gm, Rik, and Rp genes were discarded as noninformative genes. Cells with less than 500 genes per cell and with a high percentage (> 10%) of mitochondrial genes were removed from downstream analyses. Following our filtering procedures, the average UMI count per cell was 11829. The median number of genes detected per cell was 3812, 3.4% mapped to mitochondrial genes. Cell cycle annotation using the cyclone method from the scran R package[4] showed that 3366 cells were in G1 phase, 869 cells were in G2M phase, and 250 cells were in S phase. Data normalization and integration were performed using the default Seurat v4 approach FindIntegrationAnchors() followed by IntegrateData(), and differentially expressed genes were determined using a logistic regression in Seurat on the non-integrated data using the FindConservedMarkers() function. Pathway based analyses were performed using the enrichR package[5]. Unsupervised clustering was performed on the significant variable genes using the 10 first PCA followed by the nonlinear dimensionality reduction technique UMAP[6]. Annotation of the data was obtained by mapping published signatures using the AddModuleScore() method of Seurat. The MolO LT-HSC signature was taken from Wilson et al., 2015[7], and the MPP2/3/4 signatures were taken from Sommerkamp et al., 2021[8]. An Excel file listing the genes in these signatures is available in supplementary table S4. For permutation testing we randomly sampled 200 sets of genes to create gene-sets of an equivalent size to our MetaFate, Fate, transcription factor and MPP3 signatures. Expression scores for each cell were computed using the AddModuleScore() function from Seurat and the correlation between signature scores and myeloid bias was computed using a Spearmans Correlation.

**Categorisation of lineage biased barcodes:**

Labelled cells were classified as differentiation inactive (98 cells ; 61 unique barcodes) if we could not detect their respective barcode in any mature cell compartments, or as erythroid-biased (143 cells ; 35 barcodes), myeloid-biased (143 cells ; 31 barcodes), or unbiased (284 cells ; 31 unique barcodes). Lineage biases were defined based on the relative abundance of the barcode across the respective compartments. More precisely, an additional normalization step per barcode is applied in each individual, thereby enabling categorization of each barcode into classes of biased output towards the analyzed cell types. Let lineage-$bias_{bc}$ represent the relative representation of barcode *B* in cell type *C,* and let $P_{bc}$ represent the proportional read abundance per barcode per cell type

$$lineage\ bias_{bc} = \frac{P_{bc}}{\sum_c P_{bc}}$$

Specifically, a barcode was classified as myeloid biased if it was found above the 75th percentile of all myeloid-$bias_{bc}$ scores. A barcode was classified as erythroid biased if it was found in below the 25th percentile of myeloid-$bias_{bc}$ scores. A barcode was classified as unbiased if it was found between the 25th and 75th percentile of myeloid-$bias_{bc}$ scores.

These specific thresholds were set using a sensitivity analysis approach that recorded the number of cells in each category, as well as the number of differentially expressed genes and log2 fold change in gene expression between barcoded subsets. In this setting we observe that stricter thresholds in lineage bias led to greater effect sizes in gene expression changes, but at the expense of having less cells in each category (**Figure S3**). Thresholds of 25% and 75% were chosen to keep a strict criteria for lineage bias, whilst also having > 100 barcoded cells within each lineage bias category (**Figure S3**)

**Defining Metabolically-Associated Genes:** Genes are classified as metabolically associated if they are found within a metabolic pathway defined in the KEGG database, or whether they map to reactome/GO pathways associated with the following terms: *'transport', 'import','Slc', 'Abc','Atp','abc', 'metabolic process','biosynthetic process','catabolic process'*. In total 3,095 genes met this definition and the list of genes can be found in supplementary table S4.

**PAGA Trajectory Analysis:** To perform trajectory analysis, a reference map was generated from a published single-cell sequencing dataset of 44,802 C-Kit+ cells[9]. Preprocessing was performed using a scanpy pipeline as performed in (Wolf et al., 2019) and trajectory inference was performed using the PAGA method implemented in scanpy[10]. Data was then imported into Seurat and visualised using the dimensionality reduction technique UMAP[6]. Data imputation was performed using the Rmagic library[11] before calculating the expression patterns of KEGG metabolic signatures using the AddModuleScore method of Seurat.

**Causal Network Analysis of Bulk RNAseq Data** To construct a causal relationship network between our key predictive genes and a discrete lineage variable we use an information theoretic method known as miic which learns graphical models from purely observational data, including the effects of unobserved latent variables[12]. Briefly, the algorithm starts from a complete graph, the method iteratively removes dispensable edges, by uncovering significant information contributions from indirect paths, and assesses edge-specific confidences from randomization of available data. The remaining edges are then oriented based on the signature of causality in observational data.

To perform miic profiling of the haemopedia RNAseq data[13] we first processed the bulk RNAseq library. Genes with log cpm values which did not have a log cpm value of at least 1 were omitted for further analysis. This led to a dataset with 17761 genes and 110 FACs sorted haematopoietic cell population. Data was normalised using the *voom* framework in limma[14]. Differential expression analysis and pathway based analyses were also performed using the limma framework with a Benjamin Hochberg correction applied to p-values to adjust the false positive rates for multiple comparisons[14]. We next took genes from the myeloid metafate signature as well as genes encoding enzymes and transporters that were (i) upregulated in the myeloid lineage, erythroid or lymphoid lineages and (ii) variably expressed in LSK hematopoietic progenitors using published datasets[9,13]. A full list of these genes is provided in table SX. The expression patterns of these 230 genes within 104 bulk RNA seq profiles from the haemopedia database, as well as a discrete lineage variable for each sample were used as inputs into the miic web server[15] run on default parameter settings: https://miic.curie.fr/. The results of the network analysis are available at: https://miic.curie.fr/job_results.php?id=1MgzD2kVPrtuYH9vlagI

**Data and script accessibility**

**Code and data availability:** All data and code are available at https://github.com/TeamPerie/Cosgrove-et-al-2022

| Antibody target | Clone | Conjugate | Manufacturer | Relevant Figure Panel | Dilution |
|---|---|---|---|---|---|
| CD45.1 | A20 | PE/BUV737 | BD Biosciences | S11 | 1:50 |
| CD45.2 | 104 | BV605 | Biolegend | S11 | 1:100 |
| Cd34 | Ram34 | E450 | Invitrogen | 4 | 1:50 |
| Ter119 | TER119 | PE-Cy7 | BD Biosciences | 1,3,4,5,S2,S9 | 1:100 |
| CD19 | 1D3 | APC-Cy7 | BD Biosciences | 1,3,4,5,S2,S9 | 1:100 |
| CD3 | ebio500A2 | PE | eBiosciences | 1,3,4,5 | 1:100 |
| CD11b | M1/70 | E450,PerCP-Cy5.5 | eBioscience | 1,3,4,5,S2,S9,S11,S12 | 1:500 |
| CD117 (C-Kit) | 2B8 | APC/APC-Cy7 | BioLegend | 1,2,3,4,S2,S7,S9, S11,S12,S14 | 1:100 |
| CD117 (C-Kit) | ACK2 | BV650 | BioLegend | | 1:100 |
| CD135 (Flt3) | A2F10 | PE | eBioscience | 2,S7 | 1:100 |
| CD135 (Flt3) | A2F10 | PE-Cy5 | Life technologies | | 1:100 |
| Sca1 | D7 | Pacific Blue /APC-Cy7 | BioLegend | 2 | 1:200 |
| Sca1 | E13-161.7 | Pacific Blue | Biolegend | 1,2,3,4,5,S7,S9,S11, S12,S14 | 1:200 |
| CD150 | TC15-12F12.2 | PE-Cy7/ PerCPCy5.5 | BioLegend | 1,2,3,4,5,S7,S9,S11, S12,S14 | 1:100 |
| Ter119 | TER119 | biotin | BD Biosciences | Ter enrichment | 1:100 |
| CD44 | IM7 | PE | BD Biosciences | 1,3,5,S2,S9,S12 | 1:100 |
| CD41 | MVVREG30 | BV510 | BD Biosciences | | 1:100 |
| CD62L | MEL-14 | PE / BV605 | Biolegend | 2,3,4,5,S11,S12 | 1:100 |
| Cd48 | HM48-1 | APC-Cy7/PerCPCy5.5 | BD Biosciences | 2,3,4,5,S7,S9,S11, | 1:100 |

| | | | | | |
|---|---|---|---|---|---|
| | | | | S12,S14 | |
| CD16/32 | 2.4G2 | FITC | BD Biosciences | 4 | 1:100 |
| Lin | CD3ε, clone 145-2C11 ; Ly-6G/Ly-6C, clone RB6-8C5; CD11b, clone M1/70; CD45R/B220, clone RA3-6B2; TER-119 | PE | Biolegend | 4 | 1:200 |
| Lin | CD3 (17A2), Ter-119 (Ter119),B220 (RA3-6B2),Gr-1 (RB8-6C5) | PE-Cy7 | Biolegend / BD biosciences | 2,S7 | 1:200 |

**Table 1: Fluorescently labelled antibodies**

| **Genotyping** | | |
|---|---|---|
| BCM transgene | MBR26 Fwd | AAGCACTTGCTCTCCCAAAGTCG |
| | CaggRev2inn | GTAACGCGGAACTCCATATATGGG |
| Rosa26 wt | MBR26 Fwd | AAGCACTTGCTCTCCCAAAGTCG |
| | MBR 26 Rev | TCCCATTTTCCTTATTTGCCCT |
| Cre transgene | oIMR1084 | GCG GTC TGG CAG TAA AAA CTA TC |
| | oIMR1085 | GTG AAA CAG CAT TGC TGT CAC TT |
| Cre WT | oIMR7338 | CTA GGC CAC AGA ATT GAA AGA TCT |
| | oIMR7339 | GTA GGT GGA AAT TCT AGC ATC ATC C |
| **DRAG DNA Tagging PCR** | | |

| | |
|---|---|
| Preamp forward | ACTCACTATAGGGAGACGCGTGTTACC |
| Preamp reverse | GACACGCTGAACTTGTGGCCGTTTA |
| M1 tag forward | ACACTCTTTCCCTACACGACGCTCTTCCGATCNNNNNNNNNNNNCCTCGAGGTCATCGAAGTATCAAG |
| Illumina forward seq (R1) | ACACTCTTTCCCTACACGACGCTCTTCCGA*T*C (*= Phosphorothioate bond) |
| M1 rev Read2 | AGTTCAGACGTGTGCTCTTCCGATC CAGCTCGACCAGGATG*G*G |
| P5 forward | AATGATACGGCGACCACCGAGATCTACACTCTTTCCCTACACGACGCTCTTCCGATC |
| P7 index rev | CAAGCAGAAGACGGCATACGAGAT**XXXXXXX**GTGACTGGAGTTCAGACGTGTGCTCTTCCGATC |
| **Primers for targeted amplification of lineage barcodes from 10X 3' libraries** | |
| Pre-amp Partial read1 F | CTACACGACGCTCTTCCGATCT |
| PreAmp + read2 R | GTGACTGGAGTTCAGACGTGTGCTCTTCCGATCTactcactatagggagacgcgtgttACC |
| Sample index PCR P5-Rd1 | AATGATACGGCGACCACCGAGATCTACACTCTTTCCCTACACGACGCTCTTCCGATCT |
| Sample index PCR P7 Sample index | caagcagaagacggcatacgagatNNNNNNNgtgactggagttcagacgtgtgctcttccgatc |
| **Primers used in Lentiviral Barcoding** | |
| top-LIB | 5'TGCTGCCGTCAACTAGAACA-3' |
| bot-LIB | 5'GATCTCGAATCAGGCGCTTA-3' |
| 4 bp plate index forward | 5'ACACTCTTTCCCTACACGACGCTCTTCCGATCTNNNNCTAGAACACTCGAGATCAG3' |
| 4 bp plate index reverse | 5'GTGACTGGAGTTCAGACGTGTGCTCTTCCGATCGATCTCGAATCAGGCGCTTA3' |
| P5 | 5'AATGATACGGCGACCACCGAGATCTACACTCTTTCCCTACACGACGCTCTTCCGATCT3' |
| P7 | 5'CAAGCAGAAGACGGCATACGAGANNNNNNNGTGACTGGAGTTCAGACGTGCT CTTCCGATC3' |

**Table 2:** Oligo Sequences Used in This Study. Complete list of the i7 indexes in table S10. All oligo's were ordered at IDT with HPLC purified grade.

**Supplementary Tables**

Table S1: Metadata for MetaFate experiments

Table S2: Differentially expressed genes between barcoded HSPCs

Table S3: Differentially expressed pathways between barcoded HSPCs

Table S4: Gene Signatures used in scRNAseq analysis

Table S5: MetaFate expression matrices

Table S5a: Metadata for metafate Seurat object

Table S5b: Normalised metafate gene expression matrix after QC and preprocessing

Table S6: Post-QC Lentiviral Barcode Matrices

Table S6a: Lentiviral barcoding count matrix for G6PD-Tg MPPs 3 weeks post transplantation

Table S6b: Lentiviral barcoding count matrix for WT MPPs 3 weeks post transplantation

Table S6c: Lentiviral barcoding count matrix for MPP transplantation study 3 weeks post-transplantation

Table S6d: Lentiviral barcoding count matrix for Slam HSPC transplantation study 12 months post-transplantation

Table S7: Input Matric for MIIC causal network analysis

### Supplementary Figures

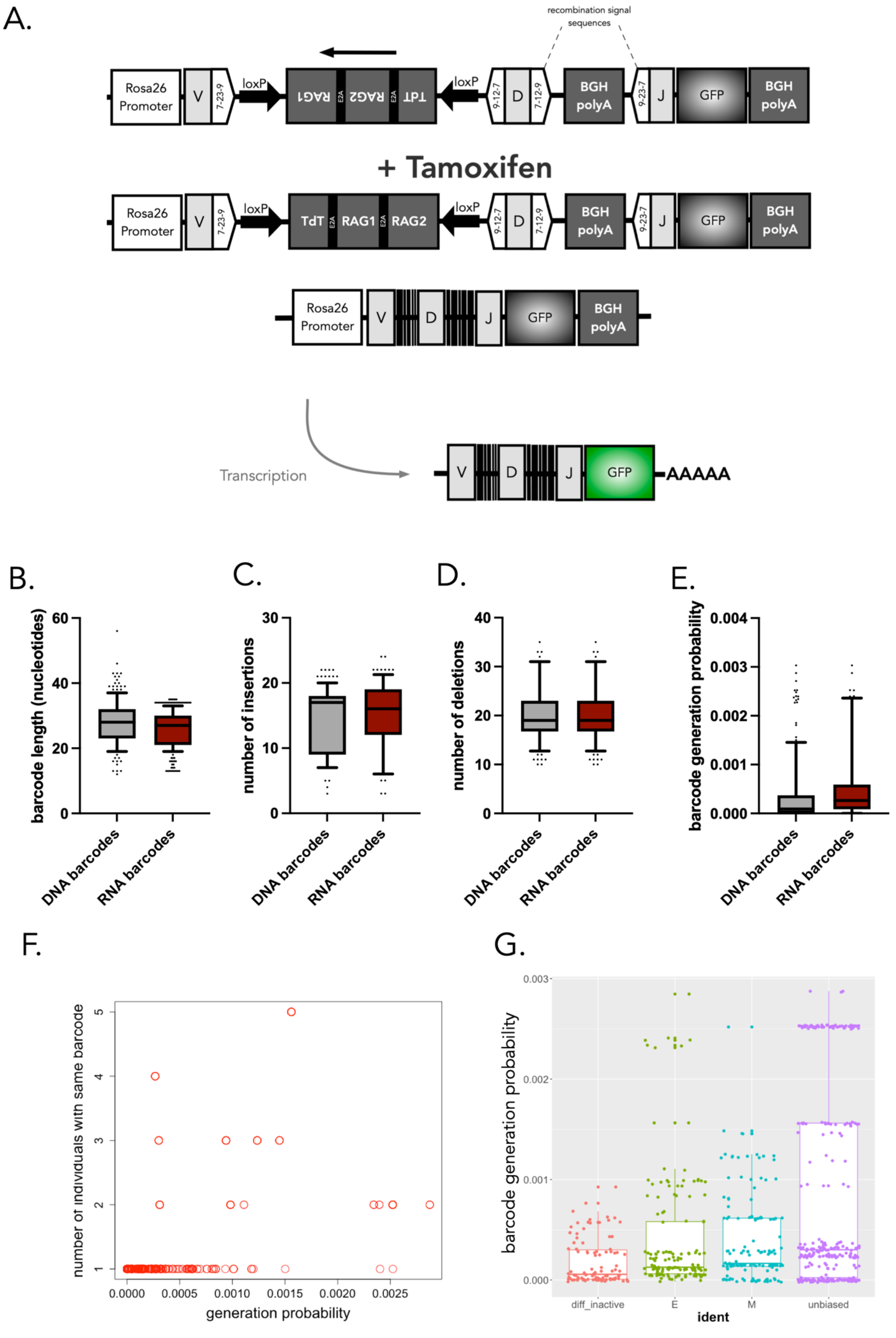
A.
recombination signal sequences
Rosa26 Promoter
V
7-23-9
loxP
RAG1
E2A
RAG2
E2A
TdT
loxP
9-12-7
D
7-12-9
BGH polyA
9-23-7
J
GFP
BGH polyA
+ Tamoxifen
Rosa26 Promoter
V
7-23-9
loxP
TdT
E2A
RAG1
E2A
RAG2
loxP
9-12-7
D
7-12-9
BGH polyA
9-23-7
J
GFP
BGH polyA
Rosa26 Promoter
V
D
J
GFP
BGH polyA
Transcription
V
D
J
GFP
AAAAA
B.
barcode length (nucleotides)
0
20
40
60
DNA barcodes
RNA barcodes
C.
number of insertions
0
10
20
30
DNA barcodes
RNA barcodes
D.
number of deletions
0
10
20
30
40
DNA barcodes
RNA barcodes
E.
barcode generation probability
0.000
0.001
0.002
0.003
0.004
DNA barcodes
RNA barcodes
F.
number of individuals with same barcode
1
2
3
4
5
0.0000
0.0005
0.0010
0.0015
0.0020
0.0025
generation probability
G.
barcode generation probability
0.000
0.001
0.002
0.003
diff_inactive
E
M
unbiased
ident

**Figure S1. Overview and QC of the DRAG barcoding technology.** (A)Description of the DRAG cassette, as inserted into the Rosa 26 locus before and after induction. DRAG recombination is induced by Cre activity and resulting barcode sequences are used for lineage tracing. The DRAG system has been designed such that upon CRE induction, a segment between two loxP sites is inverted, leading to the expression of both the RAG1 and 2 enzymes and Terminal deoxynucleotidyl transferase (TdT). Upon such expression, recognition of recombination signal sequences (RSSs) within the DRAG cassette by the RAG1/2 complex leads to recombination of the synthetic V-, D- and J-segments, with diversity being generated both by nucleotide deletion and TdT-mediated N-addition. Notably, as the RAG/TdT cassette and RSSs are spliced out during this recombination step, further recombination of the DRAG locus is prevented, and any generated VDJ sequence is thus stable over time. Finally, recombination of the DRAG locus results in the removal of a BGH polyA site that precludes GFP expression in the native DRAG configuration, allowing one to identify barcode$^{+}$ cells by flow cytometry or imaging. To allow in situ barcode generation, DRAG mice were crossed with ubiquitously expressed tamoxifen-dependent Cre (RosaCre-ER$^{TM}$) mice, to obtain the heterozygous RosaCre$^{+/-}$ DRAG$^{+/-}$ mice used in all experiments. (B) Length in nucleotide of DRAG barcodes detected from RNA or DNA (C) Numbers of insertions in DNA and RNA retrieved DRAG barcodes (D) Numbers of deletions in DNA and RNA retrieved DRAG barcodes (E) Distributions of barcode generation probabilities for RNA and DNA barcodes (F) A dotplot showing the relationship between repeat use barcodes (barcodes that occur in more than 1 mouse), and their barcode generation probability. (G) The distribution of predicted barcode generation probabilities across different classes of lineage-biased barcode categories. Data from n = 5 mice, same experiments as figure 1.

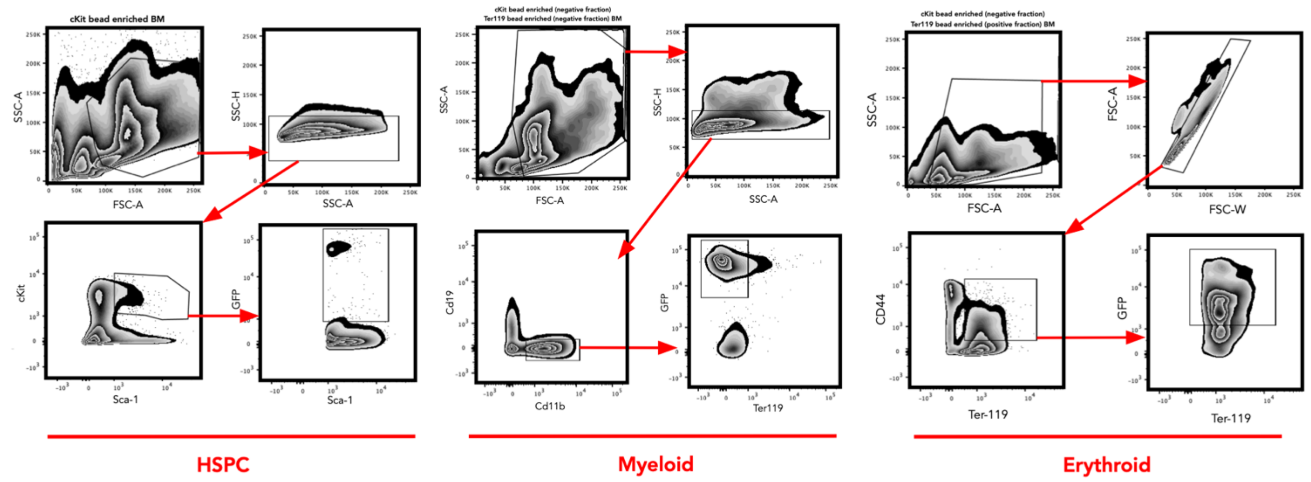


**Figure S2. Gating strategy for MetaFate experiment.** HSPCs are classified as cKit+ Sca1+ GFP+, myeloid cells are classified as CD19- Ter119- and Cd11b+ GFP+ , and nucleated erythroid cells are classified as CD19- Cd11b- and Ter119+ Cd44+ GFP+.

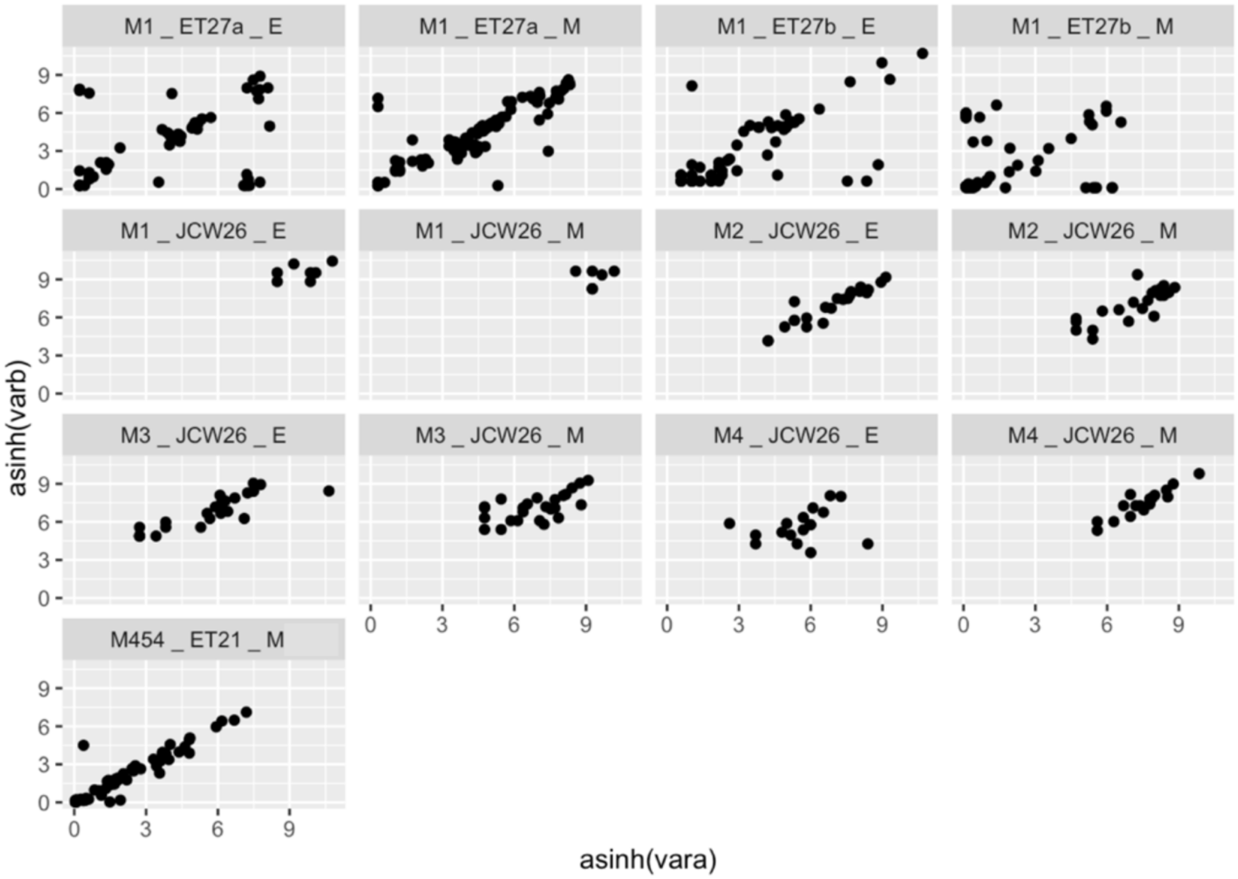


**Figure S3:** Consistency of DNA barcodes between technical duplicates. Each plot is a distinct sample, and each point represents 1 barcode**.** Data represents normalised and hyperbolic arcsin transformed read counts and each panel represents an individual sample with the sample name shown above the plot labelled as mouse_experiment code_cell type. **E corresponding to erythrocytes and M to myeloid cells.**

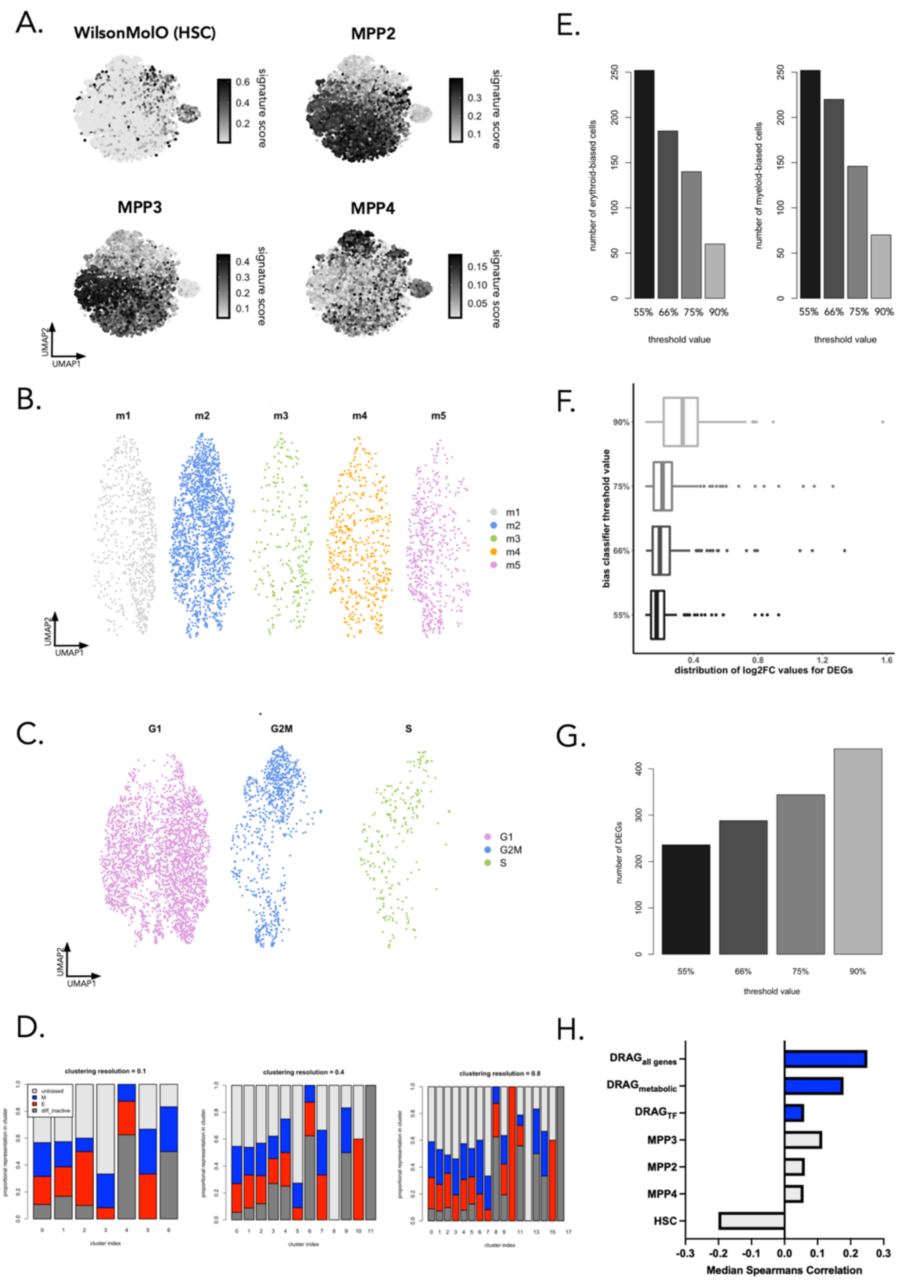


**Figure S4. MetaFate data QC. Show QC plots for 10X and for DNA barcodes.** (A) gene expression signatures for HSC (from Wilson et al[7]) and MPP subsets (from Sommerkamp et al[8] and Pietras et al.[16]) overlaid onto the UMAP representation of the data. These signatures were used to positioning cell subsets in Figure 1.B. (B) the distribution of cells from each biological replicate (individual mouse) onto the reference UMAP embedding of the data. (C ) distribution of cells in G1/G2M/S cell cycle phases in our data as determined using the cyclone method[17] implemented in the scran package. (D) The distribution of E/M/unbiased/differentiation inactive barcoded cells across unsupervised clusters for different clustering resolutions (0.1,0.4 and 0.8). This analysis was performed using Seurat's default implementation of the Louvain clustering method. (E) The number of cells belonging to each lineage bias class across a range of different lineage-bias threshold values (F) The distribution of log2fc changes in gene expression for differentially expressed genes using different lineage bias classifier threshold values. (G) The number of DEGs detected for the myeloid biased barcoded cells across a range of lineage-bias classifier values. (H) 4-fold validation of the metafate signature generation pipeline. In this analysis we partitioned barcoded cells into 4 folds that were iteratively used for training and testing. Signatures were obtained by performing differential expression between barcoded progenitor subsets and significantly genes upregulated in myeloid-biased cells were grouped into one of 3 signatures (DRAG-allgenes, DRAG-metabolic, and DRAG-TF). For each iteration (200 in total), the Spearman's correlation between gene signature and barcode derived myeloid bias was assessed and compared against published signatures from Wilson et al[7], and Pietras et al[16].

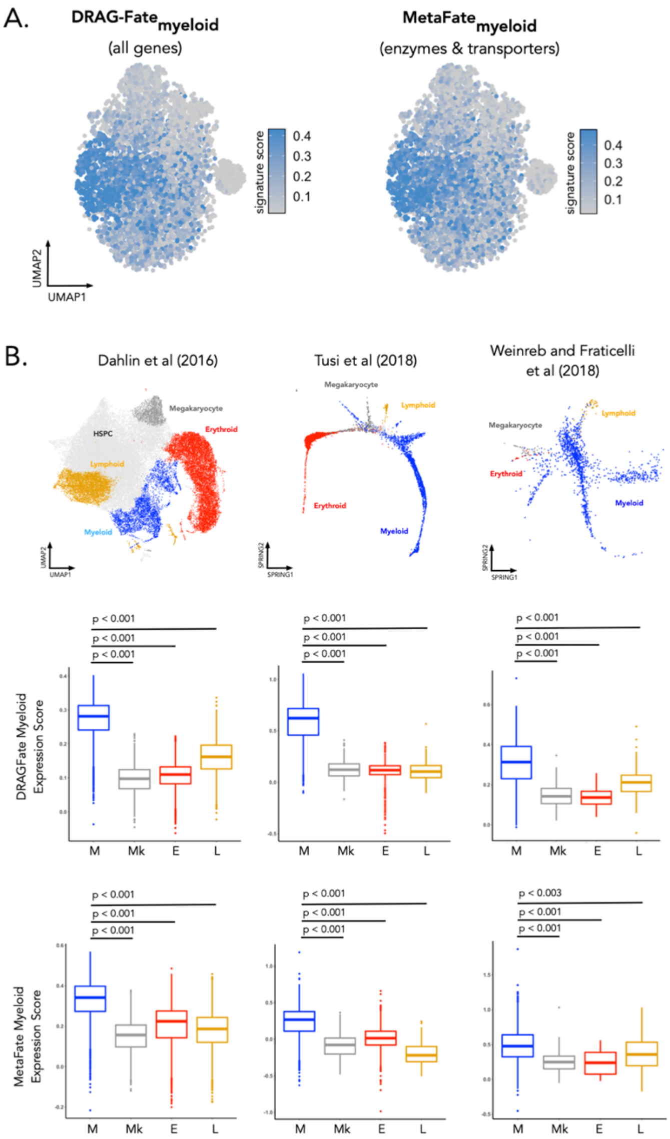


**Figure S5. Expression of the Fate-myeloid and metafate-myeloid signatures on previously published scRNAseq datasets.**(A) All genes upregulated in myeloid-biased barcoded cells compared to erythroid and differentiation inactive-barcoded cells form a gene-signature called DRAGFate-Myeloid. The subset of genes from the DRAGFate-myeloid signature relating to cellular metabolism form the MetaFate-myeloid signature. The signature score corresponds to the average expression values of these gene sets for each cell and is projected onto the UMAP visualisation of the data. (B) Benchmarking of signatures on published dataset. Here we show the expression of the DRAGFate myeloid and MetaFate myeloid signatures on published scRNAseq datasets. In these analyses we used cell-type definitions provided in the original article and signature scores were calculated using the AddModuleScore() function in Seurat. Dahlin et al (left) comprises 44,802 cKit+ and cKit+ Sca1+ hematopoietic progenitors[9]. Cell clustering and supervised assignment of cluster identity were taken from Wolf et al[10]. The dataset from Tusi et al (middle) comprises 4,763 cKit+ progenitors[18]. To annotate this dataset we performed unsupervised clustering of the data and supervised annotation using lineage-specific markers provided in supplementary table 1 of the original article[18]. Weinreb and Fraticelli et al.[19] (right) comprises 28,249 cKit+ and cKit+ Sca1+ progenitors that were lentivirally barcoded and cultured for 2 days in vitro. In this analysis cells were classified as M, Mk, E or L biased on whether the majority of cells within that clone are found within the lymphoid, myeloid, megakaryocyte and lymphoid clusters defined in the original publication. Cells that were undifferentiated or that could not be assigned to a lineage in this manner were excluded from further analysis, leaving a total of 1,518 barcoded cells for final comparisons. For all statistical tests we use a pairwise Mann-Whitney test corrected for multiple comparisons using the Benjamini-Hochberg method. M = myeloid (blue) ; Mk = Megakaryocyte (grey) ; E = erythroid (red) ; L = lymphoid (orange).

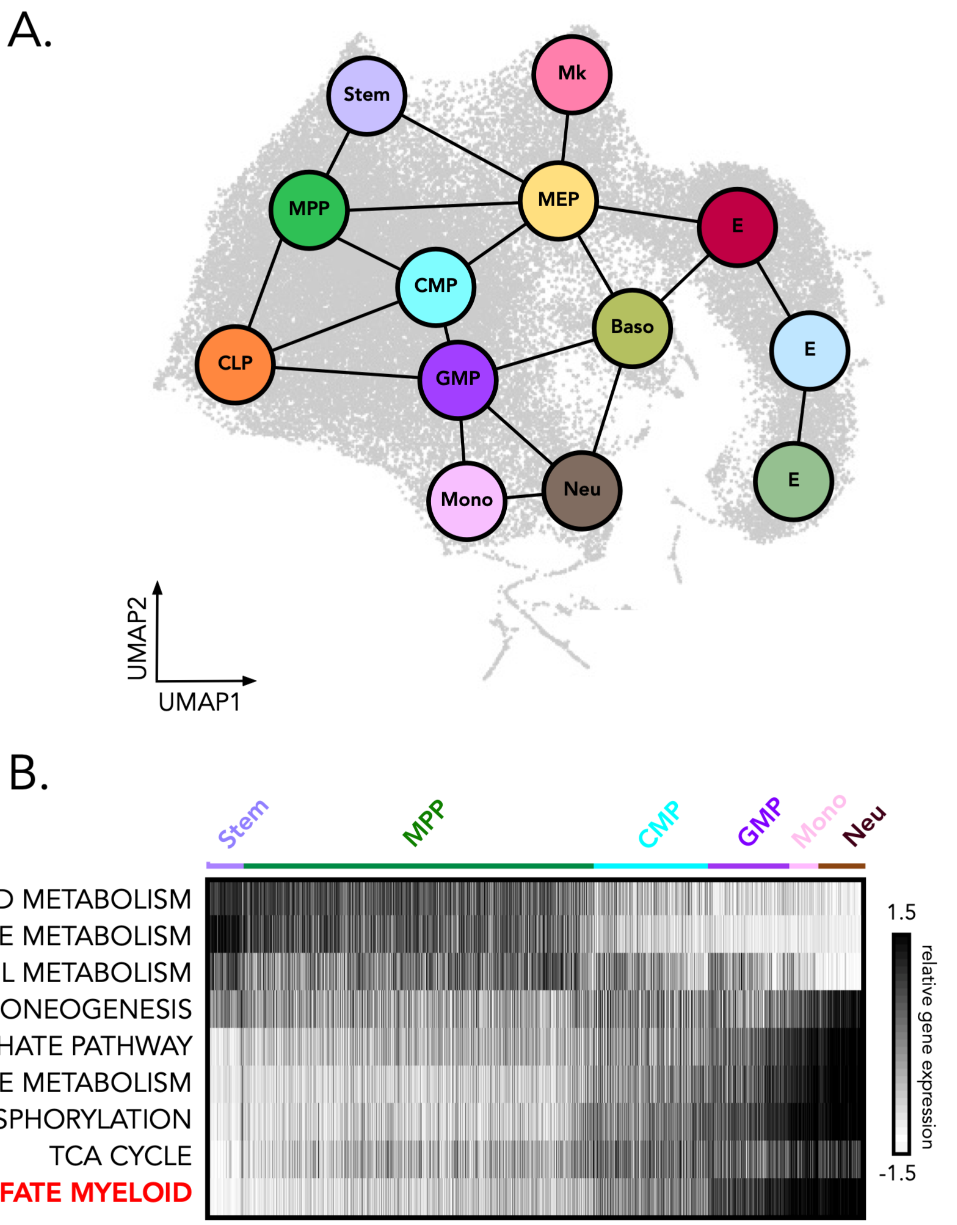


**Figure S6. The dynamics of metabolically associated gene expression patterns during myelopoiesis.** (A) A partition graph-based model of hematopoiesis generated on the scRNAseq dataset of cKit+ and cKit+ Sca1+ bone marrow cells (Dahlin et al (2018)) using the PAGA algorithm[10]. Preprocessing of the data was performed as described in Wolf et al.[10]. In this model each grey dot represents a single cell, each coloured circle represents a cell state (obtained by unsupervised clustering of gene-expression data), and edges between clusters represent putative transitions between cell states. (B) Dynamics of metabolic pathway gene expression signatures along developmental trajectories inferred by the graph based differentiation model. Each vertical line represents a single cell and the color bar at the top represents the cell states (colored circles) depicted in panel A. Pathway gene-sets were taken from the KEGG database with the exception of MetaFate myeloid and signature scores were calculated as the mean expression value across all genes in the pathway minus the mean expression values taken from randomly sampled gene-sets of an equivalent size.

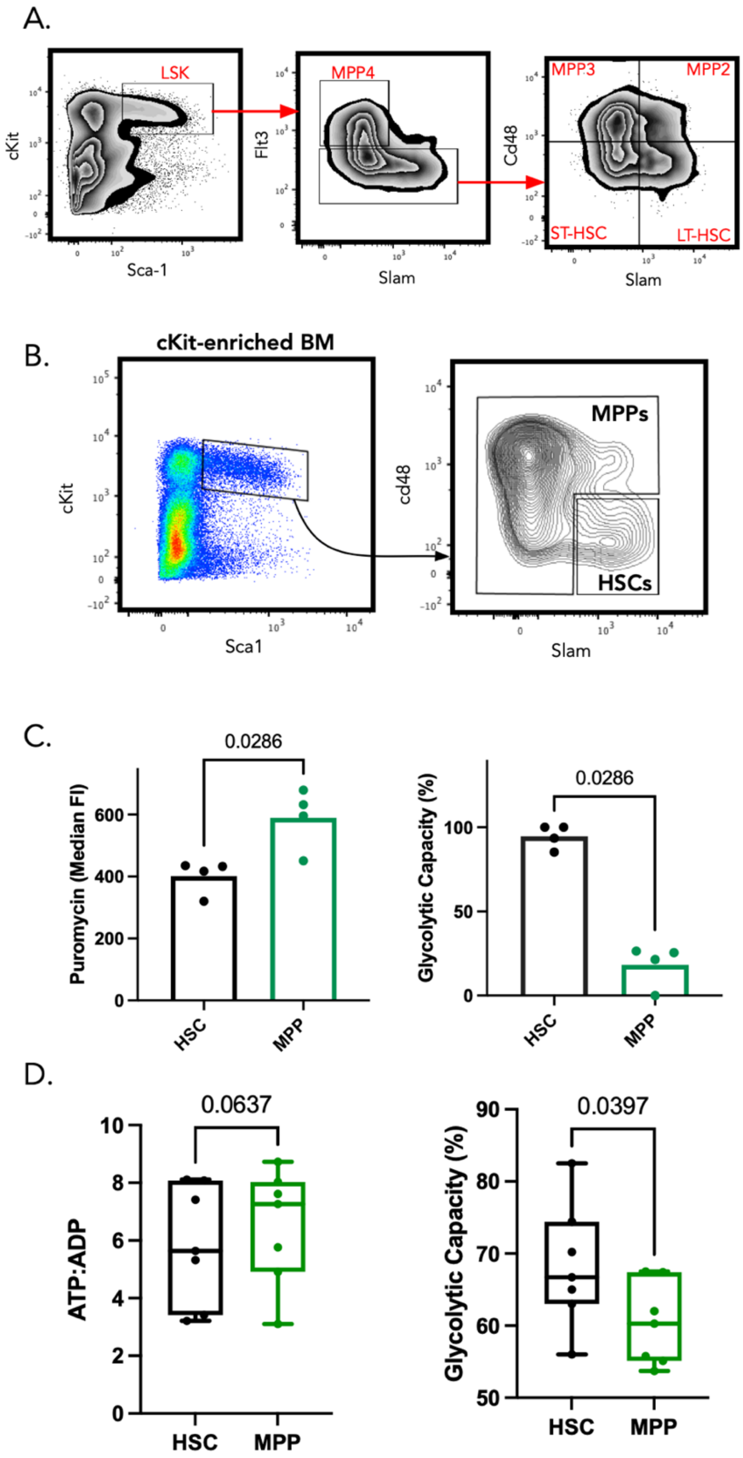


**Figure S7. Gating Strategy to define MPP subpopulations and benchmarking and validation of metabolic profiling strategy** (A) Gating strategy to define MPP2/3/4 as well as ST-HSC and LT-HSC for panel in figure 2.b. (B) Gating strategy to compare the metabolic properties of HSCs and MPPs using SCENITH and SPICE-Met (C) SCENITH profiling of HSPCs gated as in Figure 1D, each point represents a unique mouse and N=4 mice. Statistical comparisons were made using a Mann-Whitney test. (D) SPICE-Met profiling of HSPCs, each point represents a unique mouse and N=7 mice. Statistical comparisons were made using a paired T-test. Data was pooled from 2 independent experiments.

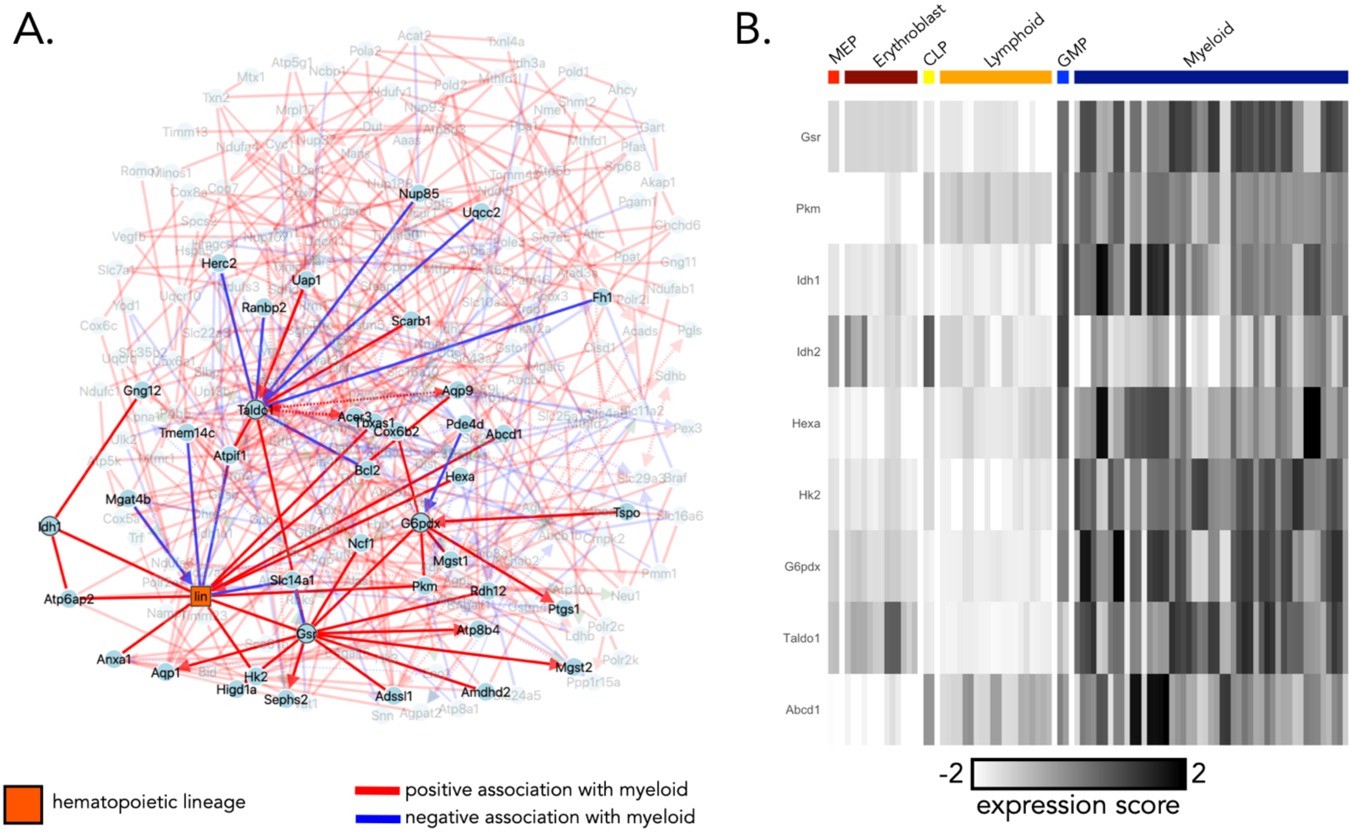


**Figure S8. MIIC network analysis to predict metabolic regulators of myelopoiesis** (A) MIIC network generated on enzyme and transporters expression patterns in bulk RNAseq patterns from the Haemopedia database. To generate an input list of enzyme and transporter genes for miic analysis we next took genes from the myeloid metafate signature as well as genes encoding enzymes and transporters that were (i) upregulated in the myeloid lineage, erythroid or lymphoid lineages and (ii) variably expressed in LSK hematopoietic progenitors using published datasets[9,13]. A full list of these genes is provided in table S7.Each circular node (blue) represents a gene. Hematopoietic cell lineage (erythroid, myeloid, and lymphoid) is represented by the square orange node and is based on which celltype the bulk RNAseq sample represents. Edges represent a predicted molecular association between genes or cell lineage with red edges highlighting genes that are positively associated with the myeloid lineage and blue lines representing genes that have a negative association with the myeloid lineage. (B) Expression patterns of enzymes/transporters that have a strong myeloid-association within the miic-generated network. Each column represents a bulk RNAseq sample taken from a FACS-sorted celltype defined in the haemopedia database.

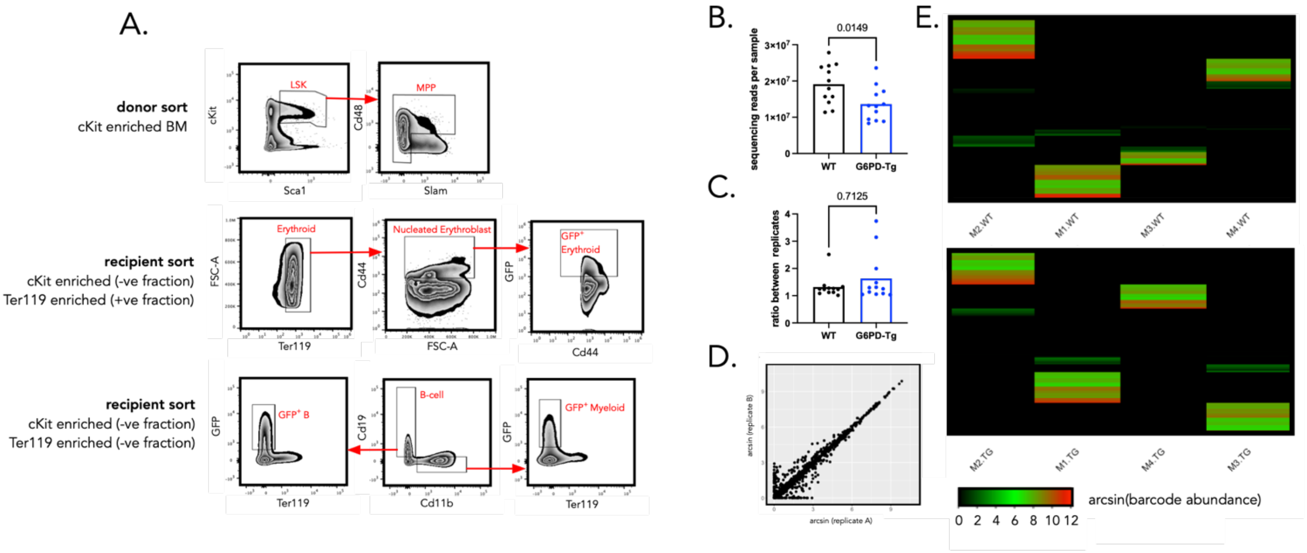


**Figure S9. Data QC for lentiviral barcoding of WT and G6PD-Tg MPPs** (A) Gating strategy to sort MPP for lentiviral barcoding experiments and to sort mature cells after transplantation. (B) The total number of sequencing reads per sample per condition. Each point represents a bulk sorted population of mature cells from one mouse. Normality was assessed using a Shapiro-Wilk test and then pairwise comparisons were made using a Students T-test (C) The ratio of barcode abundances across technical replicates in WT and G6PD-Tg conditions, each point represents a bulk sorted population of cells in one mouse. Normality was assessed using a Shapiro-Wilk test and then pairwise comparisons were made using either a Mann-Whitney test. (D) Representative scatter plot showing the consistency of read counts across technical replicates. Each point represents a unique barcode. Data are transformed using the hyperbolic arcsin function. For WT samples, the Pearson Correlation between technical replicates was of 0.89 +/-0.29 per samples and for the G6PD-tg group 0.98 +/- 0.05. (E) Heatmaps showing the distribution of barcode abundance across mice after data QC and filtering to ensure that we do not have significant numbers of barcodes shared between mice injected with the same batch, these shared barcodes are more likely to label more than 1 cell. Top heatmap represents WT samples, and bottom heatmap represents G6PD-Tg heatmaps. Data are hyperbolic arcsin transformed reads.

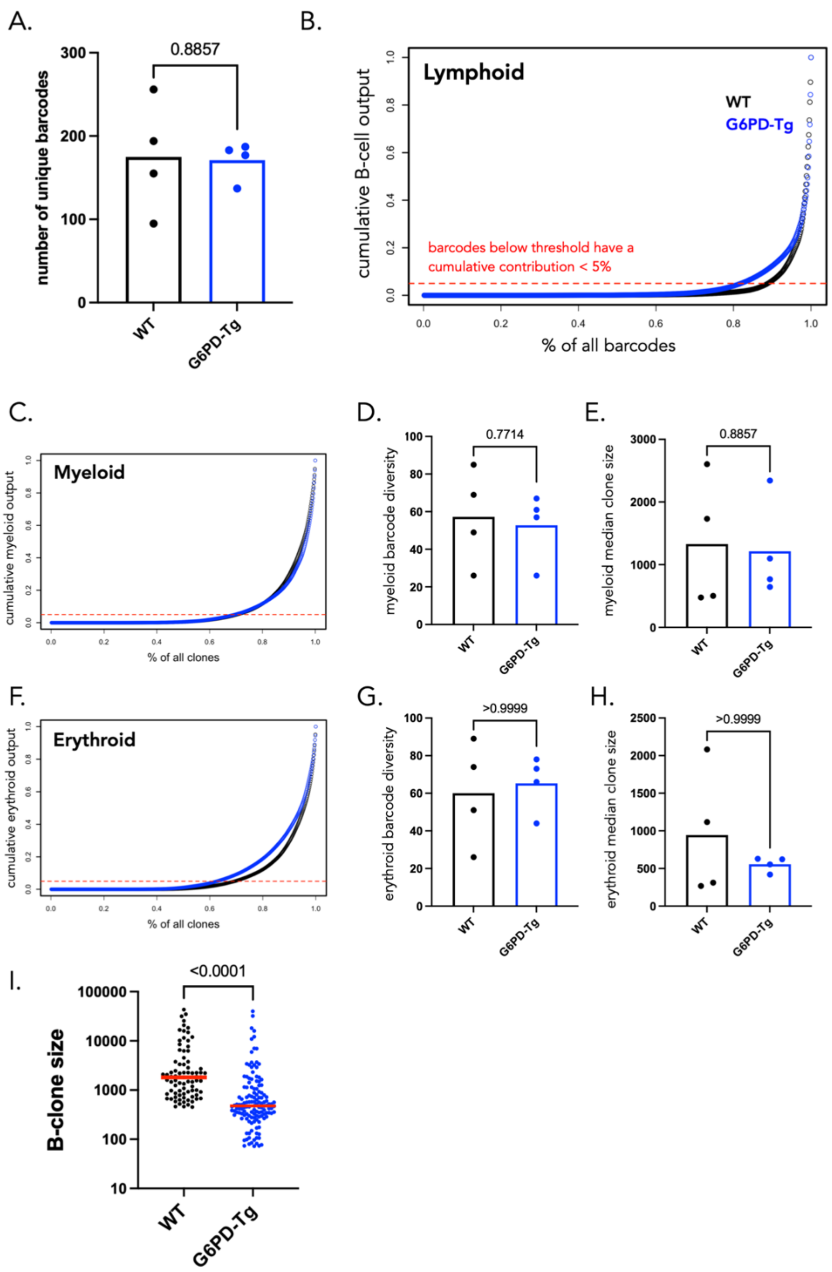


**Figure S10. Supporting analyses for lentiviral barcoding of WT and G6PD-Tg MPPs** (A) The total number of unique barcodes retrieved from WT recipient mice transplanted with WT (black) and G6PD-Tg (blue) MPPs. Each point represents 1 mouse and N = 4 mice per condition. Stats? (B) Cumulative distribution showing the abundance of each barcode in the B-cell lineage. The top *n* barcodes that contribute to 95% of all read counts for the B-lineage are classified as B-cell producing for subsequent analyses. Each point on this plot represents a distinct barcode. (C) Same as B for the Myeloid lineage. (D) The number of unique barcodes producing myeloid cells in WT recipient mice transplanted with WT (black) and G6PD-Tg (blue) MPPs. Statistical comparisons were made using a Mann-Whitney test and each point represents a unique mouse. N = 4 mice (E) The median clone size of all myeloid producing barcodes per mouse across WT and G6PD-Tg conditions. Statistical comparisons were made using a Mann-Whitney test and each point represents a unique mouse. N = 4 mice (F) Same as B for the erythroid lineage. (G) The number of unique barcodes producing erythroid cells in WT and G6PD-Tg conditions. Statistical comparisons were made using a Mann-Whitney test and each point represents a unique mouse.. (H) The median clone size of all erythroid producing barcodes per mouse across WT and G6PD-Tg conditions. Statistical comparisons were made using a Mann-Whitney test and each point represents a unique mouse. N = 4 mice (I) Distribution of clone sizes for B-cell producing barcodes across WT and G6PD-Tg conditions. Data pooled from 4 mice per condition. Each point represents a unique lineage barcode (81 WT barcodes, 138 G6PDtg barcodes). Pairwise comparisons were made using a Mann-Whitney test.

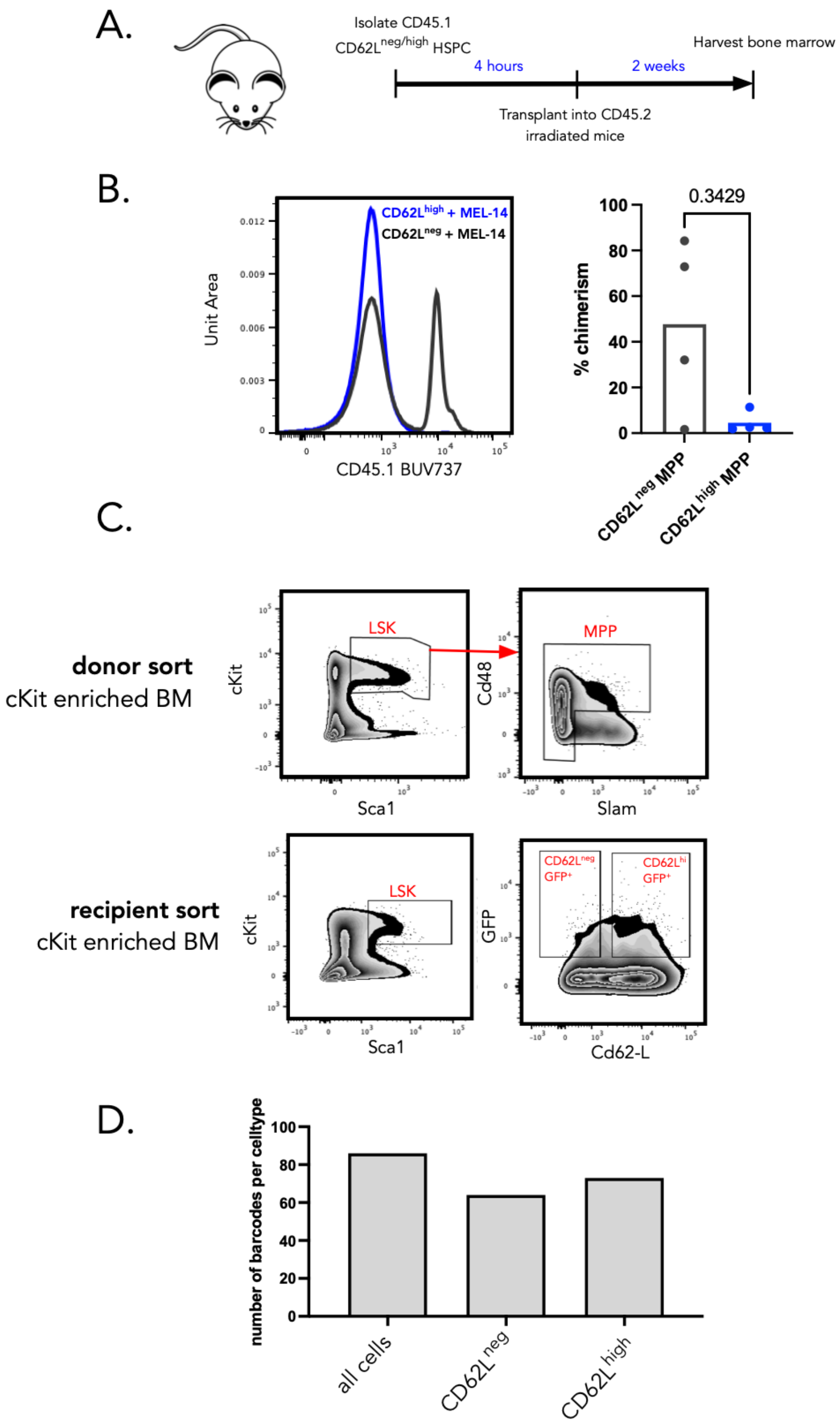


**Figure S11. Anti-CD62L (Clone MEL-14) impairs the engraftment of CD62L-expressing HSPCs** (A) Overview of the experimental timeline for the transplantation of CD62L$^{neg}$ and $^{high}$ MPPs (B) % Chimerism in cKit- bone marrow cells for the CD62L$^{neg/high}$ transplanted HSPCs. N = 4 mice per condition. (C) Gating strategy to purify MPP subpopulations pre and post transplantation. (D) Barcode diversity, as the number of unique barcode, across all cell types and for the CD62L$^{neg/high}$ subpopulations. Data pooled across all mice (n = 4)

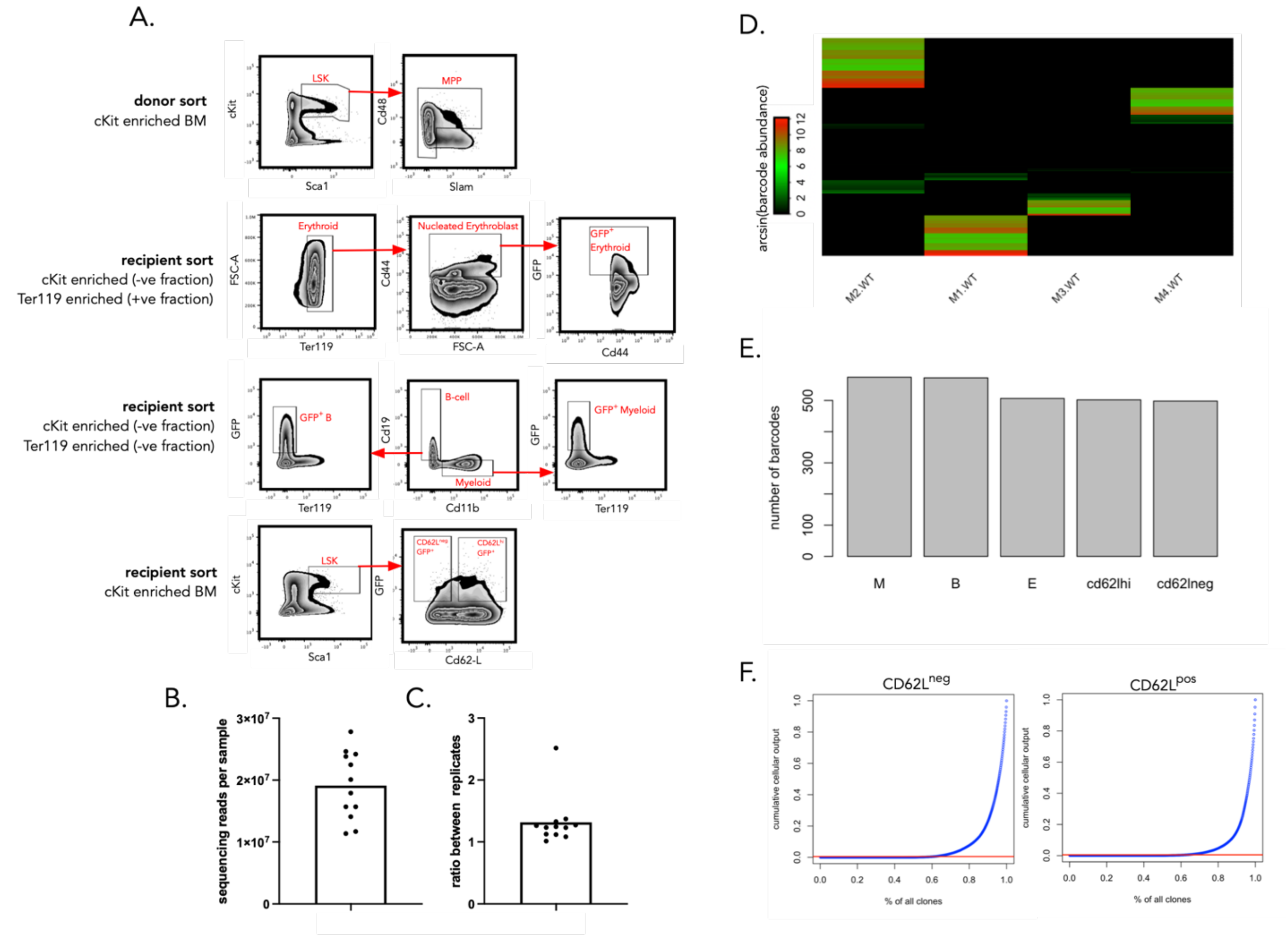


**Figure S12. lentiviral barcoding of CD62L$^{neg/hi}$ MPPs at 3 weeks post-transplantation** (A) overview of the flow cytometry gating strategy used to purify cells for lentiviral barcoding analysis of figure 5. (B) the number of sequencing reads for each sample in the barcoding analysis. Each point represents a bulk sorted cell population. N = 4 mice (C) The ratio between technical replicates where each point represents a bulk sorted cell population (D) Heatmaps showing the distribution of barcode abundance in reads across mice after data QC and filtering to ensure that we do not have significant numbers of barcodes shared between mice injected with the same batch, these shared barcodes are more likely to label more than 1 cell. Data are hyperbolic arcsin transformed reads. (E) Barcode diversity, as the number of unique barcodes, in QC processed samples pooled from 4 mice. (F) Thresholding strategy to define CD62L$^{neg/high}$ barcodes. Each point represents a single barcode and the red line represents a threshold value of 0.5%.

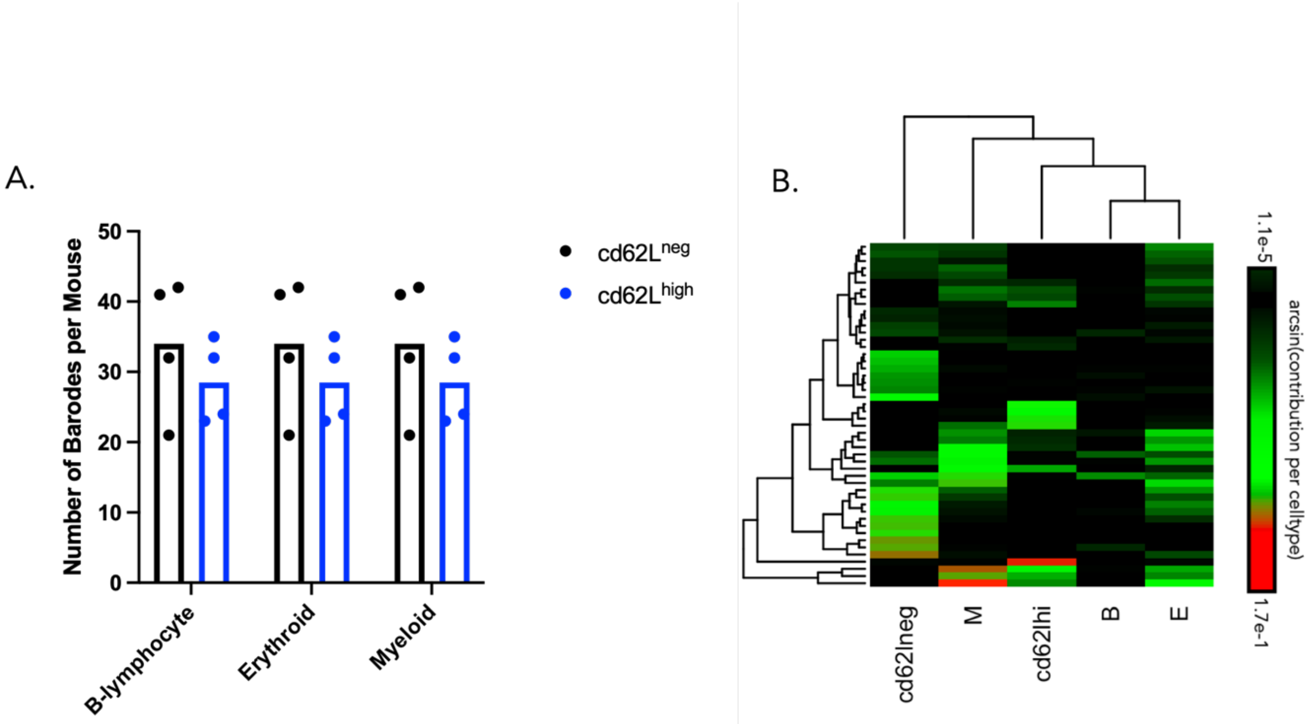


**Figure S13. lentiviral barcoding of CD62L$^{neg/hi}$ MPPs at 3 weeks post-transplantation** (A) Barcode diversity scores in myeloid (M), erythroid (E) and B-lymphocyte (B) lineages for CD62L$^{neg/high}$ MPPs (black/blue points respectively). No statistically significant differences were observed between CD62L$^{neg/high}$ conditions for any lineage. Each point represents a single mouse and statistical comparisons were made using a paired Wilcoxon-Signed Rank test. N = 4 mice (B) Unsupervised clustering and heatmap visualisation of CD62L$^{neg/high}$ barcodes across all samples. Data are pooled from 4 mice.

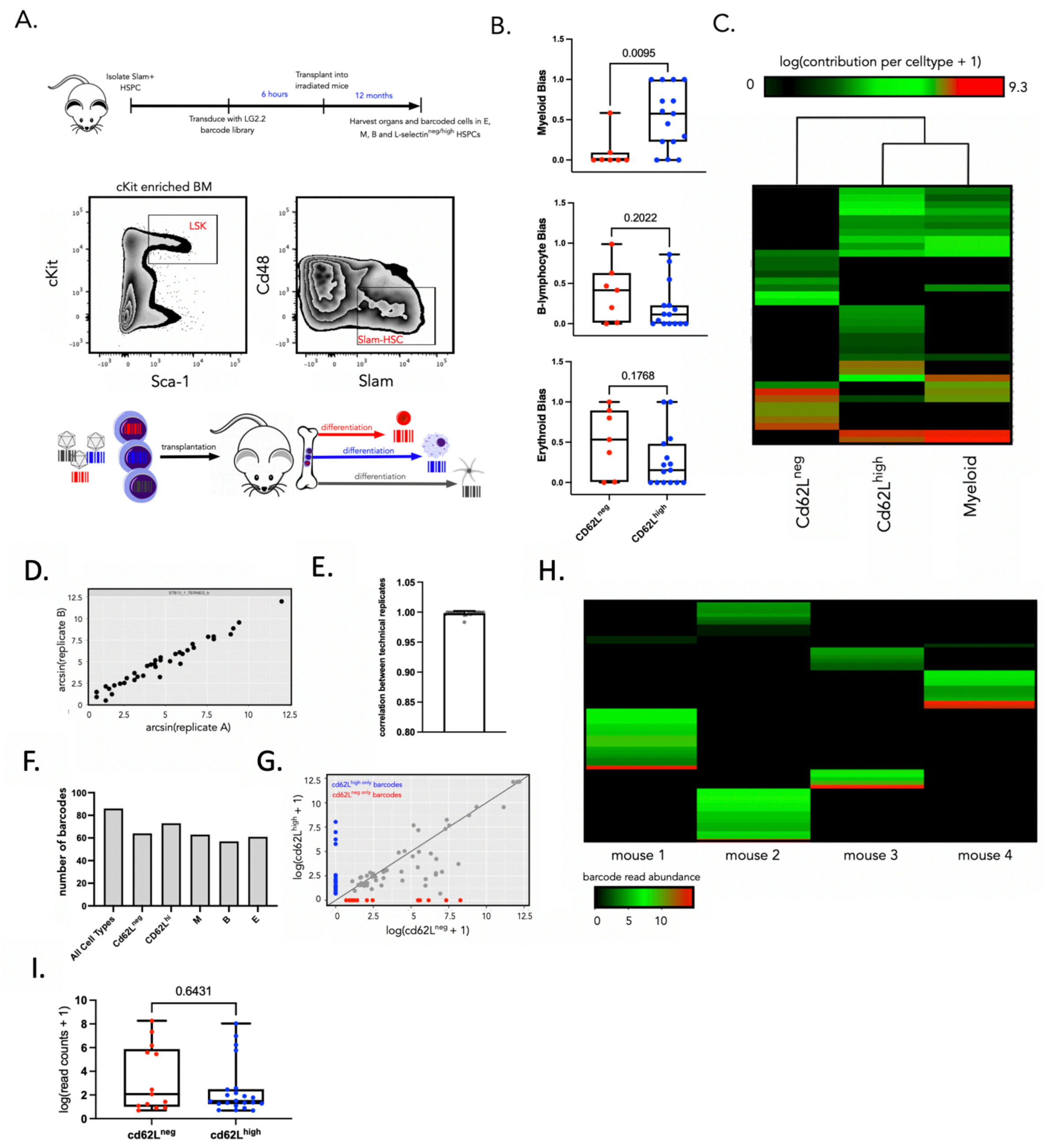


**Figure S14. The CD62L$^{hi}$ MPP compartment reconstitutes the myeloid compartment following Slam HSC bone marrow transplantation** (A) cKit+ Sca1+ CD150+ Cd48- HSPCs were purified from donor mice by FACS and infected with the LG2.2 lentiviral barcoding library. Transduced cells were transplanted into 4 sublethally irradiated (6Gy) recipient mice and 12 months later CD62L$^{neg/hi}$ MPPs, CD19$^{+}$ B cells, CD44$^{+}$ Ter119$^{+}$ erythrocytes, and CD11b myeloid cells from the bone marrow and were processed for targeted sequencing of lineage barcodes. (B) Lineage-biases of CD62L$^{neg\ only}$ or CD62L$^{hi\ only}$ MPPs for the myeloid, erythroid and B-lymphocyte lineages. Data is pooled from 4 mice, and each point represents a unique barcode. Statistical significance was assessed using a Mann-Whitney test (C) Unsupervised clustering and heatmap visualisation of barcodes that occur in either CD62L$^{neg\ only}$ or CD62L$^{hi\ only}$ MPPs. (D) An example scatter plot showing the consistency of technical replicates – each point represents a distinct barcode (E) a barplot showing the Pearsons correlation of barcode abundances across technical replicates for all samples in our dataset. Each point represents a FACs sorted bulk population of cells. (F) The number of unique barcodes detected per cell type in the lentiviral barcoding dataset. Data pooled from 4 mice (G) log barcode abundance in CD62L$^{neg}$ and CD62L$^{hi}$ MPPs. CD62L$^{shared}$ ,CD62L$^{neg\ only}$ and CD62L$^{hi\ only}$ barcodes are highlighted in grey,red and blue respectively. Data pooled from 4 mice. (H) Heatmap to show the frequency of repeat-use barcodes across mice. N = 4 mice. (I) log distribution of read counts for CD62L$^{neg\ only}$ and CD62L$^{hi\ only}$ barcodes. Statistical significance was assessed using a Mann-Whitney test.

### References for Supplementary Material


1. Eisele, A. S. *et al.* Erythropoietin directly remodels the clonal composition of murine hematopoietic multipotent progenitor cells. *eLife* **11**, e66922 (2022).
2. Faircloth, B. C. & Glenn, T. C. Not All Sequence Tags Are Created Equal: Designing and Validating Sequence Identification Tags Robust to Indels. *PLoS ONE* **7**, e42543 (2012).
3. Naik, S. H. *et al.* Diverse and heritable lineage imprinting of early haematopoietic progenitors. *Nature* **496**, 229–232 (2013).
4. Lun, A. T. L., McCarthy, D. J. & Marioni, J. C. A step-by-step workflow for low-level analysis of single-cell RNA-seq data with Bioconductor. *F1000Research* **5**, 2122 (2016).
5. Kuleshov, M. V. *et al.* Enrichr: a comprehensive gene set enrichment analysis web server 2016 update. *Nucleic Acids Res.* **44**, W90–W97 (2016).
6. McInnes, L., Healy, J. & Melville, J. UMAP: Uniform Manifold Approximation and Projection for Dimension Reduction. *ArXiv180203426 Cs Stat* (2018).
7. Wilson, N. K. *et al.* Combined Single-Cell Functional and Gene Expression Analysis Resolves Heterogeneity within Stem Cell Populations. *Cell Stem Cell* **16**, 712–724 (2015).
8. Sommerkamp, P. *et al.* Mouse multipotent progenitor 5 cells are located at the interphase between hematopoietic stem and progenitor cells. *Blood* **137**, 3218–3224 (2021).
9. Dahlin, J. S. *et al.* A single-cell hematopoietic landscape resolves 8 lineage trajectories and defects in Kit mutant mice. *Blood* **131**, e1–e11 (2018).
10. Wolf, F. A. *et al.* PAGA: graph abstraction reconciles clustering with trajectory inference through a topology preserving map of single cells. *Genome Biol.* **20**, 59 (2019).
11. Dijk, D. van *et al.* Recovering Gene Interactions from Single-Cell Data Using Data Diffusion. *Cell* **174**, 716-729.e27 (2018).

12. Verny, L., Sella, N., Affeldt, S., Singh, P. P. & Isambert, H. Learning causal networks with latent variables from multivariate information in genomic data. *PLOS Comput. Biol.* **13**, e1005662 (2017).
13. Choi, J. *et al.* Haemopedia RNA-seq: a database of gene expression during haematopoiesis in mice and humans. *Nucleic Acids Res.* **47**, D780–D785 (2019).
14. Ritchie, M. E. *et al.* limma powers differential expression analyses for RNA-sequencing and microarray studies. *Nucleic Acids Res.* **43**, e47–e47 (2015).
15. Sella, N., Verny, L., Uguzzoni, G., Affeldt, S. & Isambert, H. MIIC online: a web server to reconstruct causal or non-causal networks from non-perturbative data. *Bioinformatics* **34**, 2311–2313 (2018).
16. Pietras, E. M. *et al.* Functionally Distinct Subsets of Lineage-Biased Multipotent Progenitors Control Blood Production in Normal and Regenerative Conditions. *Cell Stem Cell* **17**, 35–46 (2015).
17. Scialdone, A. *et al.* Computational assignment of cell-cycle stage from single-cell transcriptome data. *Methods San Diego Calif* **85**, 54–61 (2015).
18. Tusi, B. K. *et al.* Population snapshots predict early haematopoietic and erythroid hierarchies. *Nature* **555**, 54–60 (2018).
19. Weinreb, C., Rodriguez-Fraticelli, A., Camargo, F. D. & Klein, A. M. Lineage tracing on transcriptional landscapes links state to fate during differentiation. *Science* **367**, (2020).
